# Mechanical signalling through collagen I regulates cholangiocyte specification and tubulogenesis during liver development

**DOI:** 10.1101/2025.01.22.633528

**Authors:** Iona G. Thelwall, Carola M. Morell, Dominika Dziedzicka, Lucia Cabriales, Andrew Hodgson, Floris J.M. Roos, Louis Elfari, Ludovic Vallier, Kevin J. Chalut

## Abstract

Cholangiocyte dysfunction accounts for a third of liver transplantations, access to which is limited by a shortage of healthy donor organs. A promising alternative is the therapeutic use of human induced pluripotent stem cell (hiPSC)-derived cholangiocytes. However, the use of hiPSCs is impeded by a lack of knowledge regarding intrahepatic cholangiocyte development, limiting the generation of fully functional cells. In this study, we generate hiPSC-derived tubular cholangiocytes using an approach based in synthetic hydrogels. These hydrogels exert control over stiffness and extracellular matrix (ECM) composition and stability, allowing us to address a critical gap in understanding cholangiocyte development. Our findings reveal that stable collagen I functionalisation, particularly on a soft substrate, enhances cholangiocyte differentiation, largely irrespective of substrate stiffness. Furthermore, high collagen I stability on a soft substrate suppresses hepatic identity whilst promoting biliary identity and duct morphogenesis. Our findings highlight the importance of collagen I mechanical signalling in regulating hepatoblast fate determination. Overall, we propose a mechanism by which the ECM modulates cholangiocyte and bile duct development and present a scalable platform for future clinical applications in the understanding and treatment of cholangiopathies.

## Introduction

The biliary tree comprises a complex network of intrahepatic and extrahepatic bile ducts, which primarily function to transport, modify and store bile from the liver to the duodenum (Brevini, Olivia C. Tysoe, and Sampaziotis 2020; Roos et al. 2022). Cholangiocytes, the epithelial cells lining these ducts, are instrumental in the modification of bile composition through absorptive and secretory processes (Maroni et al. 2015; Boyer j. and Bloomer 1974). Cholangiopathies, diseases of the cholangiocytes, lead to significant morbidity and are a major indication for liver transplantation in both paediatric (70%) and adult (33%) cases (Sampaziotis, Muraro, et al. 2021). However, liver transplantation is restricted by a lack of healthy donor organs, highlighting the urgent demand for alternative cellular therapies targeting cholangiopathies. Despite this, primary cells remain difficult to obtain in large quantities due to limited availability of primary tissue (Dhawan et al. 2020). A promising alternative is the derivation of cholangiocyte-like-cells (CLCs) from an inexhaustible source of human induced pluripotent stem cells (hiPSCs). Despite the development of multiple protocols for generating CLCs to date (Sampaziotis, C.-P. Segeritz, and Vallier 2015; Sampaziotis, Brito, et al. 2015; Sampaziotis, De Brito, et al. 2017; M. Ogawa, S. Ogawa, et al. 2015; M. Ogawa, Jiang, et al. 2021; Dianat et al. 2014; De Assuncao et al. 2015; Matsui et al. 2019; Fiorotto et al. 2018), therapeutic applications are hampered by low cell yields and a lack of functionality such as tubulogenesis (Sampaziotis, C.-P. Segeritz, and Vallier 2015; Sampaziotis, Brito, et al. 2015). Enhancing the yield and fidelity of these protocols requires a profound understanding of the developmental processes and physiological contexts that facilitate the production of fully-functional adult cholangiocytes *in vitro*.

It has been established that intrahepatic cholangiocytes originate from bipotent hepatoblast progenitors in a finely tuned developmental process (Hannan, C. P. Segeritz, et al. 2013), influenced by graded expression of biochemical signals across the lobular portal-central axis. This spatially orchestrated signalling leads to cholangiocyte differentiation at ramifications of the portal vein, coinciding with hepatocyte differentiation in the main liver parenchyma (Gordillo, Evans, and Gouon-Evans 2015; Ober and Frédéric P Lemaigre 2018; Si-Tayeb, Frédéric P. Lemaigre, and Duncan 2010; Kaylan et al. 2018; Li et al. 2018; Pauklin and Vallier 2015; Clotman, Jacquemin, et al. 2005; Clotman and Frédéric P. Lemaigre 2006; Decaens et al. 2008; Tanimizu and Miyajima 2004; Kodama et al. 2004; Zong et al. 2009). Interestingly, it was recently suggested that large collagen I deposits form in the prenatal portal triad (Jović et al. 2018), suggesting a co-operative influence of the cell niche, and particularly extracellular matrix (ECM), alongside established biochemical signals in directing cholangiocyte differentiation.

Regional variations in extracellular matrix (ECM) composition are crucial for determining cell fate and differentiation linked to ageing (Segel et al. 2019; Ge et al. 2020), development (Villeneuve et al. 2024) and disease (Koester et al. 2021; Bansaccal et al. 2023). Several studies have established a role of mechanosensitive transcriptional co-activator Yes-associated protein (YAP) (Dupont 2016) in the regulation of cholangiocyte-specific differentiation cues (Kim, So, and Shin 2023; D. H. Lee et al. 2016; L. Yang et al. 2017; Russell and Camargo 2022) and hepatocyte dedifferentiation (Blackford et al. 2023). Most studies have focused on correlating niche stiffness with cholangiocyte differentiation (Kaylan et al. 2018; Cozzolino et al. 2016; Mittal et al. 2016; Desai et al. 2016). However, emerging evidence suggests that cholangiocyte differentiation potential and phenotype are influenced by synergistic interactions between matrix composition and rigidity (Blackford et al. 2023; Kourouklis, Kaylan, and Underhill 2016; Monckton et al. 2022). Of interest, ligand-binding has been shown to adjust the stiffness-dependent threshold of YAP/TAZ (S. Lee, Alice E. Stanton, et al. 2019a; Cosgrove et al. 2016; Alice E. Stanton, Tong, and F. Yang 2019); therefore it is likely that a combined influence of substrate stiffness and matrix composition influences hepatobiliary differentiation.

In this study, we built upon our previously established polyacrylamide hydrogels (Labouesse et al. 2021; Mulas et al. 2020) to develop a tuneable synthetic hydrogel system, allowing for precise adjustment of ECM stiffness and composition. We employed this system, called StemBond hydrogels, to investigate the impact of the cell microenvironment on cholangiocyte differentiation and found that collagen I-coated hydrogels, in a largely stiffness independent manner, significantly promote cholangiocyte differentiation. This is evidenced by enhanced biliary markers and duct morphogenesis, coupled with a reduction in hepatic markers, suggesting that collagen I plays a vital role in guiding hepatoblast fate and subsequent differentiation. These findings highlight the role of the ECM, particularly collagen I, in liver development and cholangiocyte differentiation. Our findings also demonstrate the potential of our hydrogel system to explore the influence of mechanotransduction on cell fate decisions, offering a new avenue for understanding and manipulating cell behaviour *in vitro*, with implications for tissue engineering and regenerative medicine.

## Results

### Localised Collagen I in Liver Development

To better understand the microenvironment in which the biliary tree develops, we initially focused on characterising the extracellular matrix (ECM) within the human foetal liver. To investigate this, we performed immunohistological analysis on foetal liver sections ranging from 6 to 21 post-conceptional weeks (PCW) (Fig. 1.A, Fig. S1A). In accordance with previous results in foetal rat (Shiojiri and Sugiyama 2004) and human (Jović et al. 2018) livers, we observed specific localisation of collagen type I (COL I) around the portal triads. These triads consist of a hepatic artery (HA), portal vein (PV), and intrahepatic bile ducts (IHBDs) expressing keratin 19 (KRT19). Notably, such specific localisation was limited to collagen I and not present in other prevalent liver ECM components such as fibronectin, laminin and collagen type IV (Supp. Fig. 1.A). We witnessed progressive collagen I accumulation and maturation of organisation commencing at 7 PCW, preceding the appearance of robust KRT19-expressing bile ducts at 11 PCW (Fig. S1B-F). This regional collagen I increase led us to hypothesise a potential role for the instruction of the specification of cholangiocytes, the primary cell of the bile ducts, by collagen I.

**Fig. 1.**
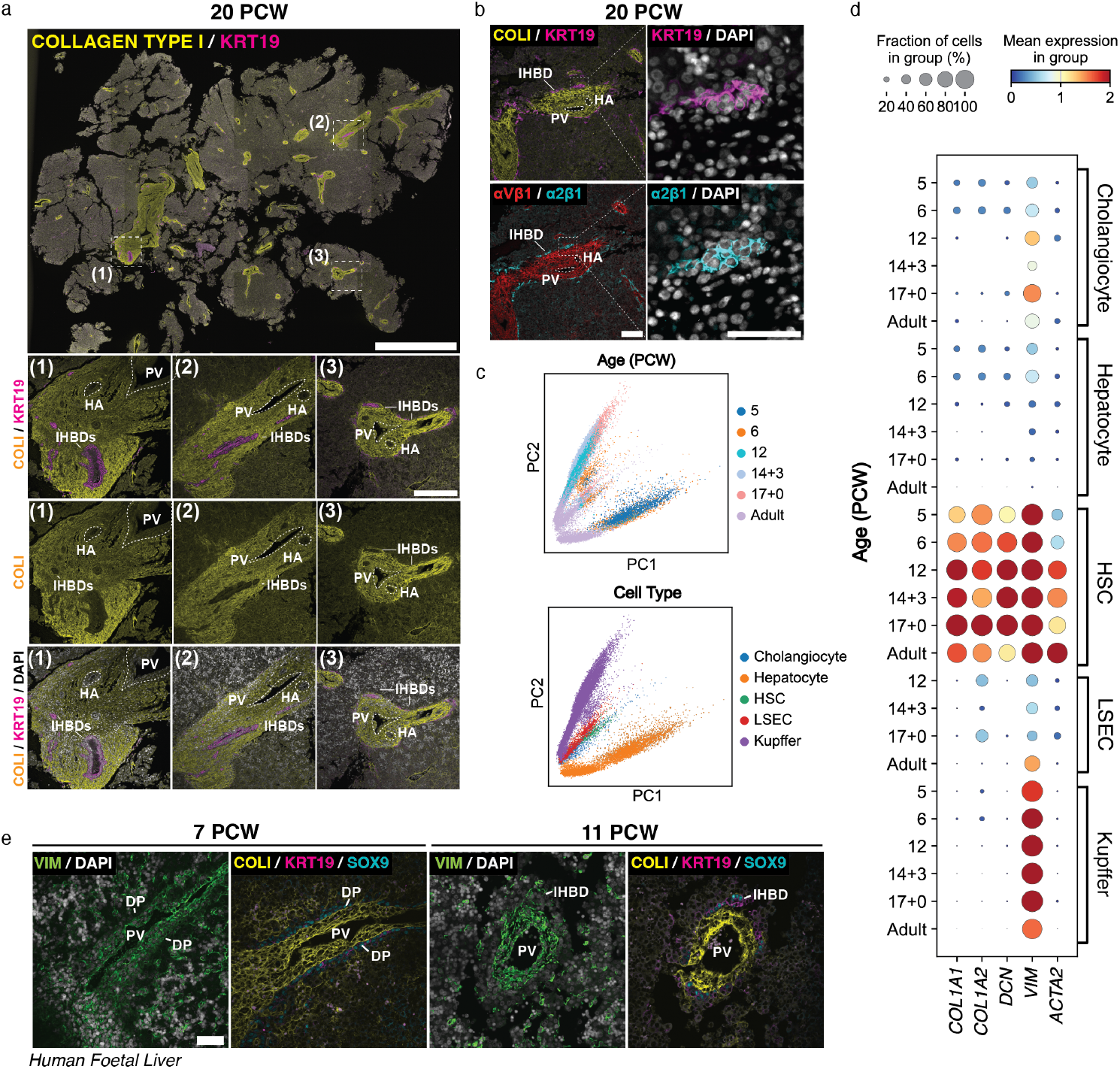
Collagen type I is specifically localised at the portal vein during *in vivo* human liver development. **(A)** Immunostaining of entire human foetal liver section at 20 post-conceptional weeks (PCW) for collagen type I (COL I, yellow) and cholangiocyte marker keratin 19 (KRT19, magenta). 10x images of regions (1), (2), and (3) are shown with and without KRT19 and DAPI (greyscale). Scale bars: 2mm and 250µm, respectively. HA: hepatic artery; PV: portal vein; IHBD: intrahepatic bile duct. **(B)** Immunostaining of human foetal liver section at 20 PCW for collagen I-binding integrin α2β1 (teal), hepatic stellate cell expressed integrin αVβ1 (red) and DAPI (greyscale) shown alongside COL I (yellow) and KRT19 (magenta) stains. Scale bars: 100µm. **(C)** PCA visualisation of all integrated single-cell transcriptomic data of foetal and adult human hepatic cells obtained via Wesley et al. [53] generated using the 10x Genomics workflow; annotation indicates cell-specific lineages (top) and PCW (bottom). **(D)** Single-cell RNA gene expression values of selected genes: hepatic stellate cell markers (decorin: *DCN*; vimentin: *VIM*; α-smooth muscle actin: *ACTA2*) and collagen I components (*COL1A1, COL1A2*) during human foetal liver development in each hepatic lineage over developmental timeframe (PCW). Dot size indicates the fraction of cells within each cell type expressing the gene and colour intensity indicates level of gene expression (‘gene expression [log-normalised, scaled counts]’) **(E)** Immunostaining of human foetal liver sections at 7 and 11 PCW for hepatic stellate cell expressed vimentin (VIM, green) and DAPI (greyscale) shown alongside COL I (yellow), KRT19 (magenta) and SOX9 (teal) stains. Scale bars: 50µm. DP: ductal plate.

Of interest, we observed a notable absence of collagen I in cholangiocytes comprising the bile ducts (Fig. 1.A), indicative of a non-parenchymal source. We speculated that hepatic stellate cells (HSCs), which are the liver myofibroblasts and collagen I producers in the fibrotic liver (Yin et al. 2013), were the main depositors of collagen I. To examine this, we conducted staining of 20 PCW sections for integrins α2β1 (ITGα2β1) and αVβ1 (ITGαVβ1) (Fig. 1.B). The former is the primary collagen I receptor, whilst the latter is a hepatic stellate cell-specific integrin implicated in TGFβ-activated procollagen I production (Han et al. 2021). Our findings reveal co-localisation of ITGα2β1 with cholangiocyte marker KRT19, whilst ITGαVβ1 was co-expressed with collagen I. This data strongly suggests that HSCs are primarily responsible for foetal collagen I production. We confirmed this via leveraging a published single-cell map of *in vivo* human liver development at developmental stages of interest (Wesley et al. 2022) (Fig. 1.C). These analyses confirm that collagen I gene expression primarily occurs in HSCs, marked by Decorin (*DCN*), Vimentin (*VIM*) and α-Smooth Muscle Actin (*ACTA2*) (Fig. 1.D).

We pursued further validation by performing immunostaining analysis of foetal liver sections at 7 and 11 PCW, focusing on the expression of non-parenchymal marker vimentin (VIM) alongside collagen I (COLI), early cholangiocyte marker SOX9 and mature cholangiocyte marker KRT19. These developmental stages were selected to explore collagen production prior to the significant accumulation observed at 20 PCW. Across both timepoints, cells surrounding the portal vein displayed co-expression of VIM and COLI, indicative of HSCs. These cells were flanked by either a ductal plate of SOX9 positive cells or intrahepatic bile ducts co-expressing SOX9 and KRT19 at 7 and 11 PCW weeks respectively (Fig. 1.E.). Interestingly, we detected VIM positive, COL I negative cells within the main liver parenchyma, which - based on the results of single cell analysis - likely represent Kupffer Cells. Taken together, our analysis reveals that collagen I organisation, a micro-architectural property associated with HSC collagen deposition, increases in the developing portal vein (Fig. S1). Thus, our analysis suggests a role of collagen I in the establishment of biliary differentiation.

### Collagen I Promotes Differentiation of hiPSCs into Cholangiocytes

In order to further explore the influence of collagen I on development, we employed an adapted protocol (Fig. 2.A) incorporating tunable polyacrylamide (pAAm) StemBond hydrogels (Labouesse et al. 2021) in the differentiation of hepatocyte and cholangiocyte-like cells (HLCs, CLCs) from hiPSCs (Sampaziotis, Brito, et al. 2015; Hannan, C. P. Segeritz, et al. 2013). Notably, StemBond hydrogels are designed to have very stable attachment of the originally seeded ECM and tunable stiffness (Labouesse et al. 2021). When seeded onto collagen I-coated StemBond (1mg/ml, rat tail), bipotent hepatoblast-like cells were capable of prolonged attachment during differentiation towards each respective fate. Of note, this was limited to collagen-coated StemBond as control conditions coated with fibronectin and laminin exhibited marked cell detachment (not shown).

**Fig. 2.**
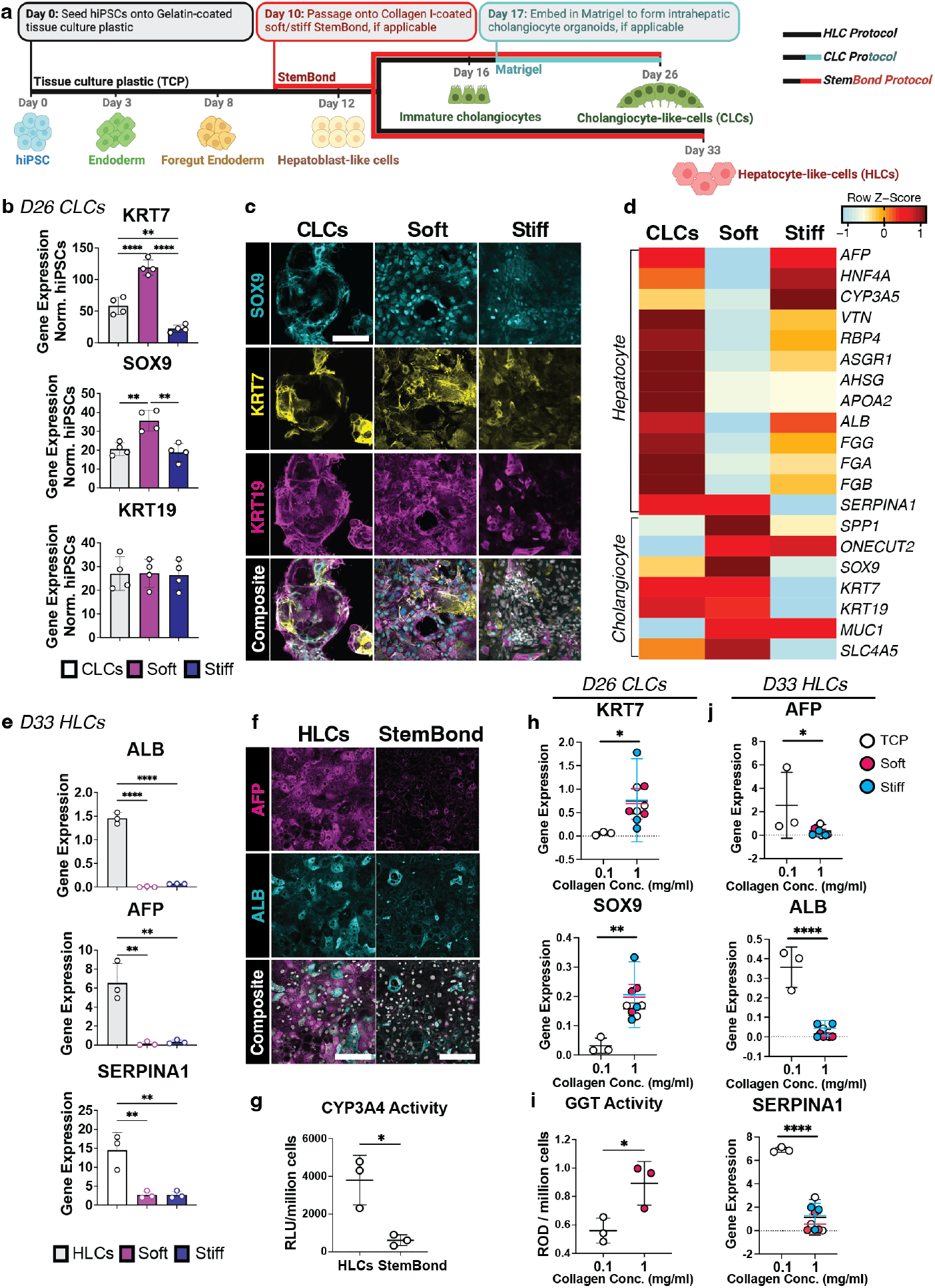
Collagen I-rich StemBond hydrogels direct hiPSC-derived hepatoblast-like cells to a biliary fate *in vitro*. **(A)** Schematic of directed differentiation from human induced pluripotent stem cells (hiPSCs) to cholangiocyte-like cells (CLCs) and hepatocyte-like cells (HLCs) *in vitro*. At day 10, when applicable, of differentiation cells are passaged onto StemBond polyacrylamide (pAAm) hydrogels coated with 1mg/ml collagen I. If not plated on StemBond, cells are embedded into Matrigel at day 17 to form intrahepatic cholangiocyte organoids (denoted as CLCs) as previously established (Sampaziotis, Brito, et al. 2015). **(B)** RT-qPCR showing mean gene expression of *KRT7, KRT19*, and *SOX9* at day 26 (*n*=4, One-Way ANOVA) relative to housekeeping genes (*PBGD, RPLP0, GAPDH*) and normalised to expression in hiPSCs. Asterisk indicates *P*-value: *≤0.05, ***≤0.001, ****≤0.0001. Error bars denote standard deviation. **(C)** Immunostaining of Day 26 cholangiocyte-like cells on soft and stiff StemBond hydrogels, compared to hiPSC-derived organoids embedded in Matrigel (CLCs). Images show expression of cholangiocyte markers KRT19 (magenta), KRT7 (yellow) and SOX9 (teal) with the nuclei stained by DAPI (greyscale). Scale bars: 100µm. **(D)** Heatmap visualisation of bulk RNA-seq (mean) for cholangiocyte and hepatocyte genes in control CLCs (*n*=2) and soft (*n*=3) and stiff (*n*=2) StemBond conditions. Heatmap colour scale displays gene expression as row Z-score. **(E)** RT-qPCR showing mean gene expression of *ALB, AFP* and *SERPINA1* (*n*=3, Kruskal-Wallis test) in day 33 hepatocyte-like cells relative to housekeeping genes (*PBGD, RPLP0, GAPDH*). Asterisk indicates *P*-value: *≤0.05, **≤0.01. Error bars denote standard deviation (SD). HLCs: gelatin TCP protocol. **(F)** Immunostaining of Day 33 HLCs on StemBond hydrogels, compared to hiPSC-derived HLCs on gelatin-coated TCP. Images show expression of hepatocyte markers ALB (teal) and AFP (magenta) with the nuclei stained by DAPI (greyscale). Scale bar: 50µm. **(G)** Cytochrome P450 3A4 (CYP3A4) activity assay for hepatocyte function (mean RLU/million cells) in day 33 HLCs (*n*=3, Unpaired T-test). Asterisk indicates *P*-value: *≤0.05. Error bars denote standard deviation. **(H)** RT-qPCR showing mean gene expression of day 26 CLCs on collagen I tissue culture plastic (TCP, white) or StemBond hydrogels (pink, soft; blue stiff) coated with high and low collagen I concentrations (1mg/ml, 100µg/ml) (*n*=3, Unpaired T-Test) relative to housekeeping genes (*PBGD, RPLP0, GAPDH*). *P*-value is indicated on graph. Error bars denote standard deviation. **(I)** Gamma-glutamyl transferase (GGT) activity assay for cholangiocyte function measured as relative optical density per million cells (mean ROD/million cells) in day 26 CLCs (*n*=3, Unpaired T-test) on collagen I tissue culture plastic (TCP, white) or soft hydrogels (pink) coated with high and low collagen I concentrations (1mg/ml, 100µg/ml). Asterisk indicates *P*-value: *≤0.05. Error bars denote standard deviation. **(J)** RT-qPCR showing mean gene expression of day 33 HLCs on collagen I TCP (white) or StemBond (pink, soft; blue stiff) coated with high and low collagen I concentrations (1mg/ml, 100µg/ml) (*n*=3, Unpaired T-Test) relative to housekeeping genes (*PBGD, RPLP0, GAPDH*). *P*-value is indicated on graph. Error bars denote standard deviation.

Given the robust attachment of hepatoblasts to collagen-I coated StemBond, we went on to investigate the potential of Collagen I-coated StemBond as a substrate for CLC and HLC differentiation. CLCs generated with the Collagen-StemBond protocol expressed cholangiocyte markers at levels comparable or superior to hiPSC-derived CLCs (Sampaziotis, Brito, et al. 2015; Sampaziotis, C.-P. Segeritz, and Vallier 2015) (Fig. 2.B-D) and akin to primary intrahepatic cholangiocyte organoid (ICOs) controls (Fig. S2.A). Moreover, we found that substrate stiffness exerted an impact on differentiation, as evidenced by enhanced cholangiocyte marker expression on soft StemBond hydrogels when compared to stiff (Fig. 2.B-D) coinciding with a notable decrease in hepatocyte marker expression compared to control CLCs (Fig. 2.D, Fig. S2.B). From these analyses, we concluded that both collagen I matrix and substrate stiffness play a role in the observed cholangiocyte specification. Gene set enrichment analysis (GSEA) provided weight to this hypothesis, as genes involved in the “collagen-containing extracellular matrix” were differentially expressed between StemBond and control CLC conditions (Fig. S2.C).

To further investigate the role of collagen I and substrate stiffness in differentiation, we cultured HLCs using the Collagen-StemBond protocol and observed reduced hepatocyte and greater cholangiocyte marker expression compared to the standard gelatin TCP protocol (Fig. 2.E-F, Fig. S2.D-E) (Hannan, C. P. Segeritz, et al. 2013). Interestingly, this analysis suggests that the suppression of hepatocyte phenotype may be partly substrate stiffness-dependent but is likely primarily driven by the culture conditions offered by StemBond regardless of stiffness. We further showed, using a cytochrome P450 3A4 (CYP3A4) assay, that culture on Collagen-StemBond significantly decreased hepatocyte functionality (Fig. 2.G).

To examine the influence of collagen I, we proceeded to differentiate CLCs and HLCs on tissue culture plastic coated with varying collagen concentrations (0.1mg/ml, 1mg/ml). We observed that greater concentrations of collagen I significantly enhanced cholangiocyte marker expression, to a similar level to StemBond (Fig. 2.H) and showed a decrease in hepatocyte markers (Fig. S2.F). A similar trend was present in gamma-glutamyl transferase (GGT) activity, with greater activity on soft Collagen-StemBond (Fig. 2.I). Furthermore, HLCs regained hepatic marker expression in decreased collagen I concentration conditions (Fig. 2.J). On the other hand, hepatocyte marker expression in CLCs was suppressed on soft StemBond versus TCP using the same collagen concentration (Fig. S2.F). Thus, some benefit appears to be conferred by the ECM stability and mechanical conditions offered by StemBond. Taken together, our analyses suggest that not only the presence but also the amount and stability of collagen I is an essential factor in directing the fate of hiPSC-derived hepatoblasts into hepatocytes and cholangiocytes. From this point forward, we restrict our 2D culture conditions to collagen I–coated soft StemBond.

### Collagen I Binding in hiPSC-derived CLCs and Primary Organoids

Given the association between collagen I and biliary differentiation, we investigated the gene expression of collagen-binding integrin subunits in CLCs cultured on soft StemBond hydrogels coated with 1mg/ml collagen I, further referred to as “StemBond”. Among these subunits, *ITGB1* emerged as the most highly expressed across two different timepoints (Days 16 and 26), with significantly higher expression compared to all other subunits (Fig. 3.A). This is notable given that ITGα2β1, the presence of which we confirmed via immunostaining (Fig. S3.A), binds to collagen I fibrils with high specificity (Jokinen et al. 2004). Additionally, immunostaining for the active configuration of ITGβ1 showed differences in protein localisation and expression between StemBond and control CLC conditions. On StemBond, there was enhanced localisation on the cell surface membrane and overall quantified intensity (Fig. 3.B-C). These data suggest increased receptor-substrate binding on collagen I-functionalised StemBond. Supporting this, gene enrichment analysis revealed upregulation of genes characteristic of matrix degradation, including *MMP12* and *MMP14*, in StemBond compared to control CLCs (Fig. S3.B), an alteration typically observed in response to integrin-mediated collagen I binding (Borrirukwanit et al. 2014). This provides further support for our hypothesis that integrin-collagen binding plays a role in CLC differentiation.

**Fig. 3.**
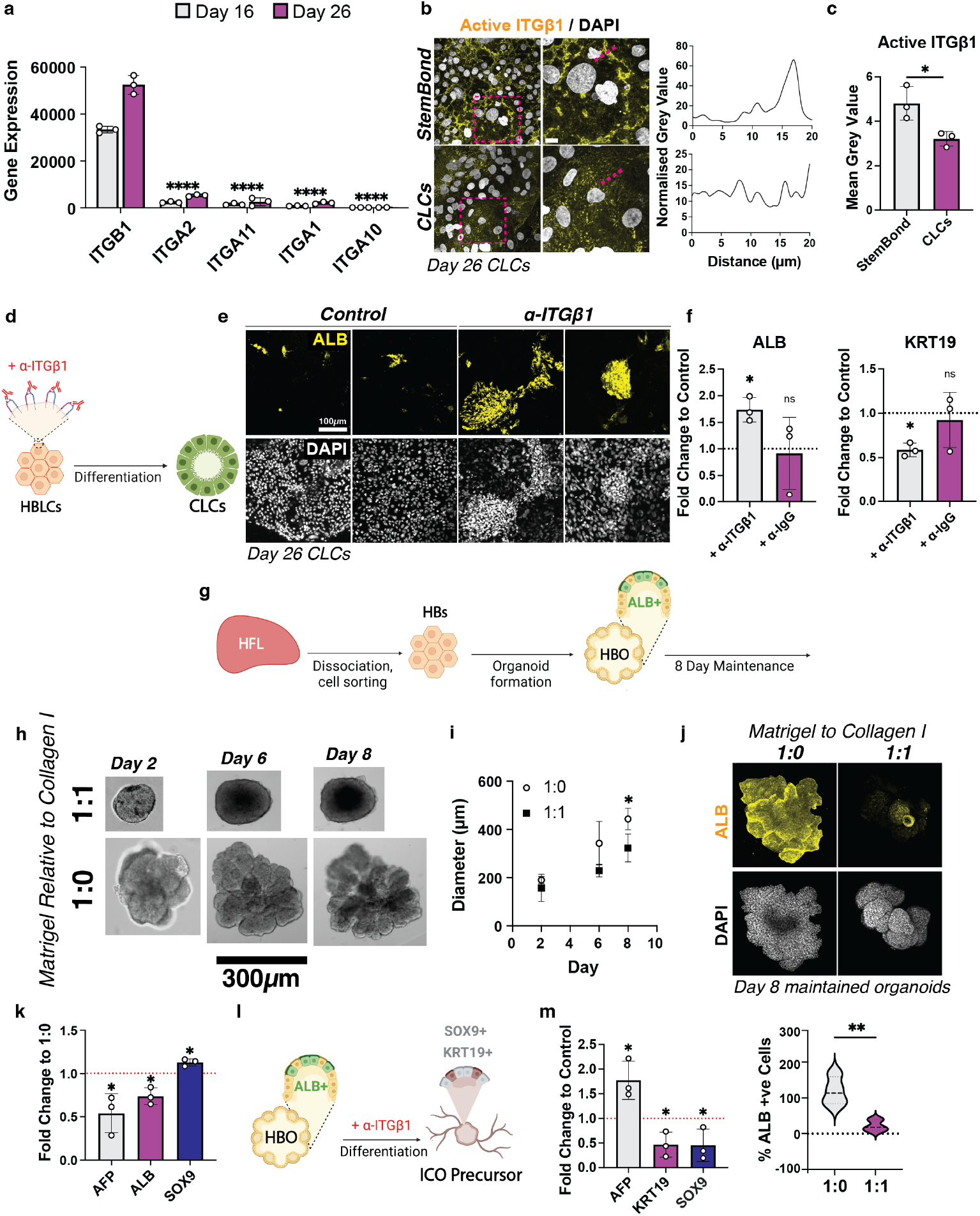
Collagen I-binding reduces hepatocyte marker expression *in vitro* as measured in hiPSC-derived hepatoblast-like cells (A-F) and primary human hepatoblast organoids (HBOs, G-N). **(A)** Bulk RNA-seq mean gene expression values indicating the expression of collagen-binding integrin subunits (*ITGB1, ITGA2, ITGA11, ITGA1, ITGA10*) in day 16 and 26 cholangiocyte-like cells (CLCs) cultured on soft collagen I-rich StemBond hydrogels (*n*=3, Multiple Unpaired T-tests). Asterisk indicates *P*-value compared to *ITGB1*: **** ≤ 0.0001. Error bars denote standard deviation. **(B)** Max projection of confocal images of day 26 StemBond and control CLCs. Cells are stained for ITGβ1 (yellow) and nuclei are stained with DAPI (greyscale). Scale bars: 25µm. Graphs show fluorescence intensity measurements along the area indicated by the red line. **(C)** Quantification of mean ITGβ1 fluorescence intensity normalised to background intensity (*n*=3, Unpaired T-test). Asterisk indicates *P*-value: * ≤ 0.05. Error bars denote standard deviation. **(D)** Schematic depicting inhibition of collagen binding in hepatoblast-like cells (HBLCs) using an ITGβ1 blocking antibody (α-ITGβ1) prior to CLC differentiation. **(E)** Max projection of confocal images of day 26 StemBond CLCs with or without α-ITGβ1. Cells are stained for ALB (yellow) and nuclei are stained with DAPI (greyscale). Scale bars: 100µm. **(F)** RT-qPCR showing mean expression of cholangiocyte (*KRT19*) and hepatocyte (*ALB*) marker genes in day 26 StemBond CLCs supplemented with α-ITGβ1 (*n*=3, Unpaired T-test). Values indicate integrin β1 blocking antibody conditions (+α-ITGβ1) and IgG controls (+α-IgG) relative to control unsupplemented conditions. Asterisk indicates *P*-value: *≤0.05, ns≥0.05. Error bars denote standard deviation. Dashed line indicates control values. **(G)** Schematic representation of the isolation of hepatoblasts (HBs) and subsequent derivation of HBOs from human foetal liver (HFL) tissue. **(H)** Bright-field images showing HBOs cultured in maintenance media and either a 1:0 or 1:1 mix of Matrigel and collagen I over 8 days, scale bar: 300µm. **(I)** Quantification of mean organoid diameter is shown (*n*=3, Unpaired T-tests). Error bars denote standard deviation. Asterisk indicates *P*-value: *≤0.05 when compared to the 1:0 control. **(J)** Max projection confocal images of maintained HBOs across culture conditions (1:0, 1:1). Organoids are stained for ALB (yellow) and nuclei are stained with DAPI (greyscale). Scale bars: 166µm. The percentage of ALB-positive cells normalised to DAPI (median ± range) is plotted (*n*=3, Unpaired T-test). Asterisk indicates *P*-value: **≤0.01. **(K)** RT-qPCR for mean expression of cholangiocyte (*SOX9*) and hepatocyte (*ALB, AFP*) marker genes in day 8 1:1 HBOs (*n*=3, Unpaired T-test). Values are displayed as fold change compared to the 1:0 control (indicated by the dashed line). Asterisk indicates *P*-value: *≤ 0.05. Error bars denote standard deviation. **(L)** Schematic of differentiation of HBOs to intrahepatic cholangiocyte precursor organoids (ICO precursor), either with or without blocking antibody α-ITGβ1. **(M)** RT-qPCR for mean expression of cholangiocyte (*KRT19, SOX9*) and hepatocyte (*AFP*) marker genes in day 8 differentiated HBOs supplemented with α-ITGβ1 (*n*=3, Unpaired T-test). Values are displayed as fold change compared to the uninhibited control (indicated by the dashed line). Asterisk indicates *P*-value: *≤0.05. Error bars denote standard deviation.

To further investigate the importance of integrins, we treated cells on StemBond with the ITGβ1 blocking antibody (αITGβ1, clone AIIB2), known to functionally inhibit ITGβ1 (Hutton et al. 2010) (Fig. 3.D). In the presence of an ITGβ1 blocking antibody, we observed an enhanced level of ALB-expressing cells, indicative of hepatocytes, whilst CLC marker KRT19 was significantly reduced, changes which weren’t present in αIgG-supplemented controls (Fig. 3.E-F). All in all, the above data indicates that collagen I binding via ITGβ1 plays a crucial role in steering hepatoblast differentiation towards a biliary fate, whilst minimising hepatocyte differentiation. The observation that collagen I binding steers hepatoblasts towards a biliary fate was further validated using primary hepatoblast organoids (HBOs), a model system capable of differentiating into either hepatic or biliary organoids, dependent on culture conditions. Utilising our established HBO protocol (Fig. 3.G), we investigated the influence of collagen I on hepatoblast differentiation through varying concentrations of Matrigel relative to a collagen I hydrogel. We maintained HBOs in a 1:1 mix of collagen I gel and Matrigel, producing a final concentration of 1mg/ml collagen I, and compared them to HBOs cultured in Matrigel alone (1:0). Over 8 days of culture, HBOs maintained in collagen conditions exhibited notable morphological differences to HBOs maintained in Matrigel alone, condensing into smaller structures characteristic of the initial stages of cholangiocyte specification (Fig. S3.C), possessing a reduced surface area and fewer projecting buds (Fig. 3.H-I). Moreover, immunostaining for hepatocyte marker ALB displayed a significant shift between conditions (Fig. 3.J), with lower ALB in the presence of excess collagen, suggesting reduced hepatocyte differentiation in comparison to control HBOs. This trend persisted at the gene expression level (Fig. 3.K), with a marked decrease in hepatocyte/hepatoblast markers (*ALB, AFP*) in the 1:1 collagen condition coupled with increased cholangiocyte marker (*SOX9*) expression. This reduction of characteristic hepatoblast features in collagen conditions suggests that collagen I influences their self-renewal capacity *in vitro*.

In order to investigate the influence of collagen I on hepatobiliary fate choice, we used a previously published protocol (Wesley et al. 2022) to facilitate the induction of biliary fate. To achieve this, we modified the culture environment through removal of small molecule inhibitor A-83-01 and addition of TGFβ, conditions previously demonstrated to induce biliary differentiation and reduce hepatic marker expression. As expected, HBOs cultured in both 1:0 and 1:1 conditions acquired branching morphologies by day 8. However, whilst uniform expression of hepatocyte marker ALB was observed across conditions, biliary marker expression appeared enhanced in the 1:1 condition (Fig. S3.D-E). Furthermore, the average branch length was significantly greater in the 1:1 condition as quantified across multiple biological replicates. Overall, these observations may indicate enhanced biliary fate choice in the 1:1 condition. To further investigate the role of integrin β1 during HBO differentiation to cholangiocytes, we proceeded to inhibit its function with the blocking antibody (Fig. 3.L,M). Gene expression analysis post-inhibition (Fig. 3.M) revealed significantly greater hepatic marker expression (*AFP*) and reduced biliary markers (*KRT19, SOX9*) when compared to control conditions, which was further validated by immunofluorescence analysis (Fig. S3.F). Taken together, these findings further indicate that collagen I concentration and integrin β1-mediated interactions direct hepatoblast fate choice and differentiation pathways, with increased collagen binding correlated with hepatocyte inhibition.

### Integrin *β*1-Mediated Collagen I Binding is Associated with YAP/TAZ Signalling

Having established a role of Collagen-StemBond in mediating hepatoblast fate choice, we next sought to clarify which signalling pathways relevant to that fate choice are activated on StemBond. To test the role of signalling in our system, we compared marker expression to that observed in two previously established protocols: the embedding of CLCs in Matrigel and the 2D differentiation of CLCs on gelatin-coated TCP (Sampaziotis, Brito, et al. 2015). Through GSEA of the KEGG and WikiPathways (WP) pathway databases, we identified genes related to PPAR signalling as one of the top five differentially expressed groups (Fig. 4.A), with *PPARA* and *PPARG* strongly downregulated on day 16 StemBond compared to the CLC control (Fig. 4.B). This was notable given that PPAR signalling acts on hepatocyte differentiation via suppression of mechanically induced YAP/TAZ signalling (Kim, So, and Shin 2023; Dupont et al. 2011), a known activator of biliary differentiation (Molina et al. 2021), which correlates with protein conjugation efficiency (S. Lee, Alice E Stanton, et al. 2019b) and inhibits hepatic specification (Noce et al. 2019). Thus, we hypothesised that YAP may be involved here in the inhibition of hepatic marker expression. We then used an existing human foetal liver scRNA-seq dataset to show elevated expression of YAP markers and target genes in developing cholangiocytes compared to hepatocytes (Fig. S4.A) (Wesley et al. 2022). We further supported this finding through immunostaining of human foetal liver sections at 7 PCW, revealing co-localisation of active YAP with cholangiocyte markers KRT19 and SOX9 (Fig. S4.B). These changes in gene expression validated previously established results, implicating YAP in cholangiocyte development L. Yang et al. 2017. Furthermore, CLCs generated via the StemBond protocol highly express YAP markers and signalling targets after 16 and 26 days (Fig. 4.C), with enhanced target expression most pronounced at day 26. The upregulation of YAP signalling at day 16 was confirmed via GSEA using published datasets of signalling pathways (Fig. 4.D) and cellular identities (Fig. 4.E). Reduced hepatic cell fate in StemBond coincided with greater expression of genes associated with YAP and associated pathways Mothers Against Decapentaplegic Homolog 2/3 (SMAD2/3) (Szeto et al. 2016) and Activator Protein 1 (AP1) (Koo et al. 2020). Immunostaining analysis corroborated this as day 16 cells on StemBond exhibited a significantly higher nuclear to cytoplasmic ratio of active YAP than those in ICO conditions (Fig. 4.F), as quantified across multiple biological replicates, with nuclear YAP coinciding with nuclear SOX9 (Fig. S4.C). The observed difference persists at day 26, with collagen I-coated TCP rescuing the reduction of nuclear YAP observed in TCP conditions (Fig. S4.D). All in all, this data shows a correlation between PPAR inhibition and YAP/TAZ activation with collagen I adherence, which results in inhibition of hepatic fate choice.

**Fig. 4.**
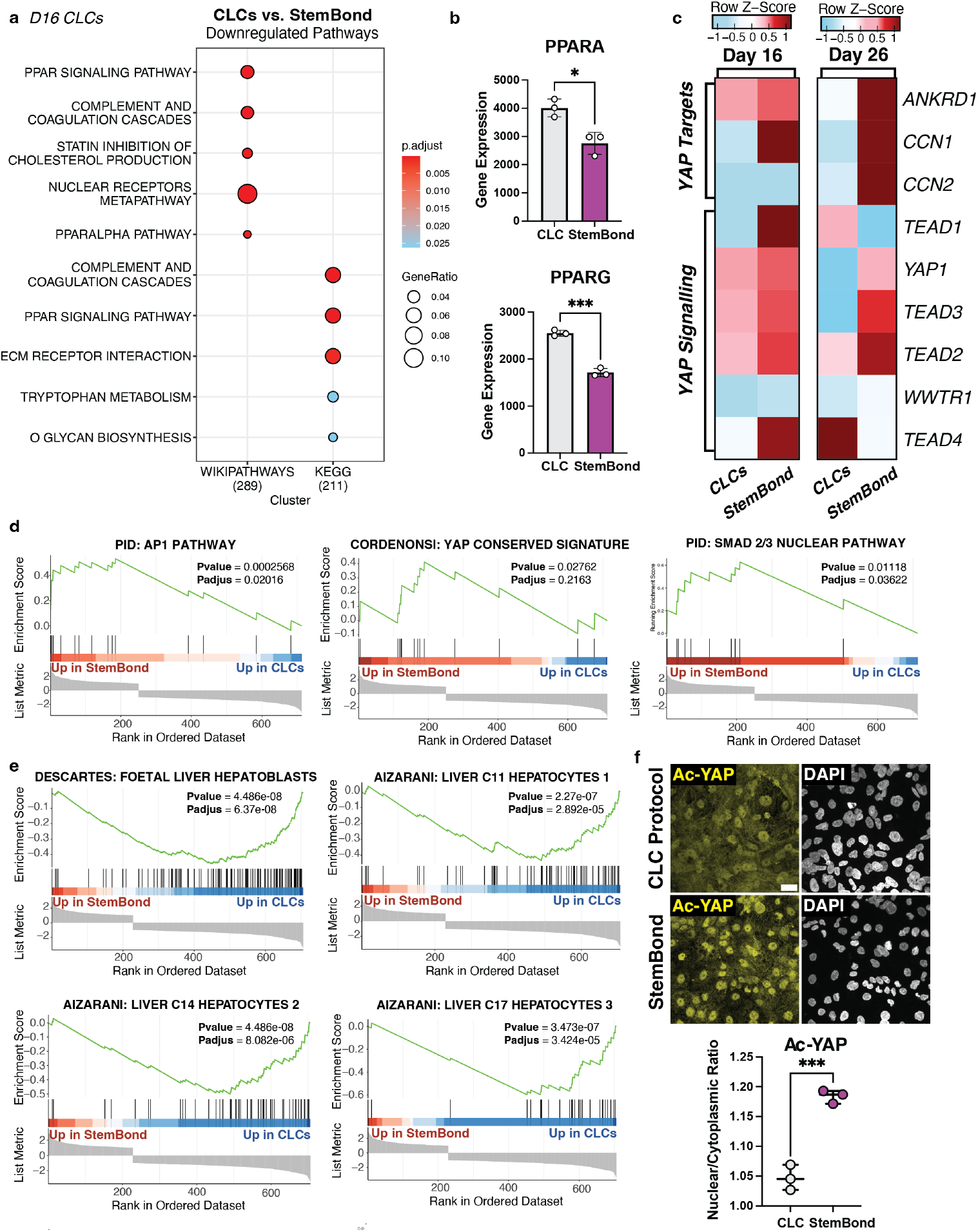
YAP/TAZ signalling in cholangiocyte-like cells (CLCs) is upregulated on collagen-rich substrates. **(A)** Dot plot of GSEA analysis (log2FC≥1, Adjusted *P*-value ≤0.05) showing the top 5 differentially down-regulated gene sets on Day 16 StemBond compared to CLC controls (indicated by line colour) from the KEGG and WikiPathways Gene Databases. Dot size indicates number of genes expressed per dataset and colour indicates adjusted *P*-value. **(B)** Bar plot representation of mean bulk RNA-seq gene expression (*n*=3, Unpaired T-test) of upstream inhibitors of YAP (*PPARA, PPARD, PPARG*) in day 16 hiPSC-derived CLCs. CLCs: hiPSC-derived intrahepatic cholangiocyte organoid protocol. Error bars denote standard deviation. **(C)** Heatmap depicting mean bulk RNA-seq expression of YAP marker genes (*YAP1, TEAD1, TEAD2, TEAD3, TEAD4*) and targets (*ANKRD1, CCN1, CCN2*) in day 16 and 26 hiPSC-derived CLCs. Colour scales indicate row Z-Score of each timepoint. CLCs: hiPSC-derived intrahepatic cholangiocyte organoid protocol. **(D-E)** Gene set enrichment analysis (GSEA) of differentially expressed genes (log2FC≥1, Adjusted *P*-value ≤0.05) between day 16 cells in StemBond and control CLC conditions. Normalised enrichment score and *P*-value are shown. Gene sets represent signalling pathways of interest **(D)** and target cell types **(E). (F)** Maximum intensity projections of confocal images staining for active YAP (Ac-YAP, yellow) and nuclei (DAPI, greyscale) in day 16 CLCs. Scale bar: 25µm. The active YAP nuclear/cytoplasmic intensity ratio is quantified across biological replicates (*n*=3, Unpaired T-test). Asterisk indicates *P*-value: ***≤0.001. StemBond; CLCs passaged onto collagen I-coated pAAm hydrogels, CLCs; Non-passaged CLCs on gelatin-coated TCP and later embedded in Matrigel as previously established (Sampaziotis, Brito, et al. 2015; Hannan, C. P. Segeritz, et al. 2013).

### Collagen I Promotes Tubulogenesis of Cholangiocytes

Intrahepatic bile duct development can be broadly categorised into two stages: first, cholangiocyte specification and second, tubulogenesis of the intrahepatic biliary tree. Therefore, we sought to investigate whether the ECM-related mechanisms we identified as crucial for fate specification also influence the process of tubulogenesis.

Building upon our differentiation protocol, we began to observe self-organisation into tubular structures, termed Tubular Cholangiocyte-like cells (Tubular CLCs), during CLC differentiation on Collagen-StemBond (Fig. 5.A,B). This tube morphogenesis was noticeably absent in CLCs cultured on TCP coated with an equivalent collagen concentration (1mg/ml). Considering the developmentally regulated properties of collagen fibrils *in vivo* (Fig. S1.D-F), we hypothesised that collagen fibril organisation may be the underlying reason for this disparity. Scanning Electron Microscopy (SEM) analysis revealed that collagen fibrils on StemBond exhibited significantly greater organisation compared to those in TCP conditions (Fig. 5.C). This enhanced organisation on StemBond suggests that collagen fibril arrangement is indispensable in the initiation of cholangiocyte tube formation.

**Fig. 5.**
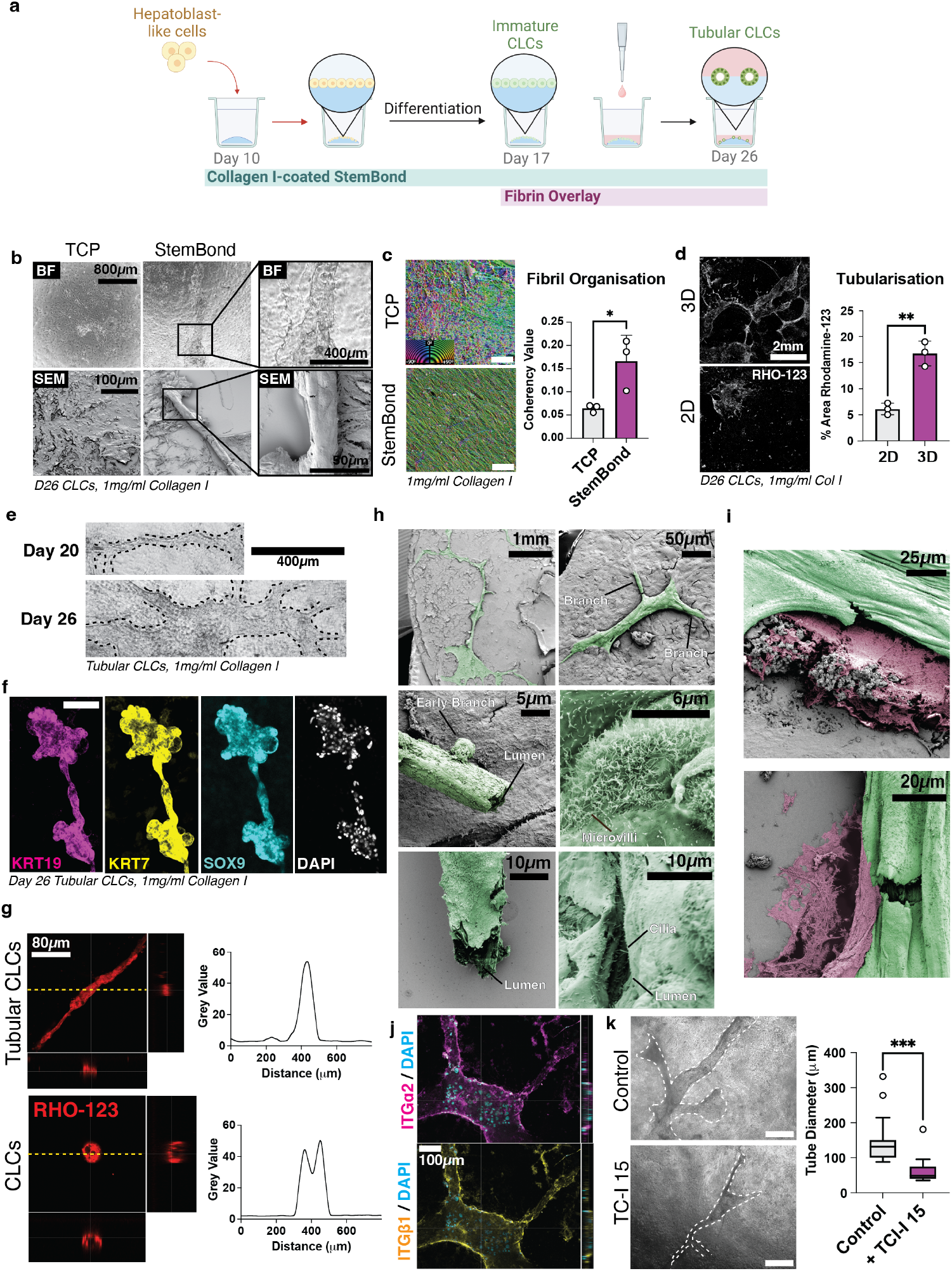
Collagen I-rich 3D sandwich gels produce tubular bile duct structures from hiPSC-derived cells *in vitro*. **(A)** Schematic representation of 3D synthetic hydrogel protocol, where day 17 cholangiocyte-like cells (CLCs) cultured on collagen I-rich StemBond polyacrylamide hydrogels are coated with a collagen I functionalised agarose fibrin overlay to form tubular CLCs. **(B)** Brightfield (BF) and Scanning Electron Microscopy (SEM) visualisation of day 26 CLCs cultured on tissue culture plastic (TCP) or StemBond hydrogels functionalised with 1mg/ml collagen I. Scale bars indicated in black. **(C)** SEM images of collagen fibrils (1mg/ml) on StemBond and TCP. Fibril orientation represented by colour (green = more aligned). Scale bar: 3µm. Fibril organisation (orientation mean coherency value) quantified across biological replicates (*n*=3, Unpaired T-test). Asterisk indicates *P*-value: *≤0.05. Error bars denote standard deviation. **(D)** Whole well images of tubularisation (stained by Rhodamine-123, RHO-123, greyscale) of day 26 CLCs cultured in 2D and 3D StemBond conditions with and without the agarose fibrin overlay. Scale, 2mm. Tubularisation was quantified across biological replicates (*n*=3, Unpaired T-test) by mean percentage area Rhodamine-123. Asterisk indicates *P*-value: **≤0.01. Error bars denote standard deviation. **(E)** Brightfield images of day 20 and 26 Tubular CLCs (outlined by dashed line). Scale bar: 400µm. **(F)** Max projection confocal images and orthogonal views of day 26 Tubular CLCs stained for KRT19 (magenta), KRT7 (yellow) and SOX9 (teal). Nuclei are co-stained with DAPI (greyscale). Scale, 150µm. **(G)** Max projection of confocal images of Tubular CLCs and CLC controls assayed for luminal Rhodamine-123 (RHO-123) internalisation (red) after 40min. Scale bar: 80µm. Dashed line represents region of quantification. Fluorescence intensity measurements (normalised to the highest value) across the dashed line are plotted. **(H-I)** False coloured SEM images of day 26 Tubular CLCs (green) **(H)** Labelled arrows show regions of interest. Scale bars indicated in black. **(I)** Collagen I fibrils are shown in pink. Scale bars: 25µm and 20µm, from top to bottom. **(J)** Orthogonal view confocal stainings of day 26 Tubular CLCs for integrin subunits α2 and β1 (ITGα2, magenta; ITGβ1, yellow). Scale bar: 200µm. Nuclei are co-stained with DAPI (cyan). Images used are of Tubular CLCs on StemBond with no agarose fibrin overlay. **(K)** Brightfield images of day 26 Tubular CLCs (outlined in white) cultured with or without ITGα2β1 inhibitor TC-I-15. Scale bar: 800µm. Plots show quantification of media tube diameter (*n*=3, Mann-Whitney test). Asterisk indicates *P*-value: ***≤0.001. Box, interquartile range (IQR); whiskers, range (minimum to maximum).

As hiPSC-derived CLCs previously required 3D culture conditions for organisation into ring-like structures (Sampaziotis, Brito, et al. 2015), we reasoned that tubularisation could be upregulated via the addition of a collagen I-functionalised agarose-fibrin hydrogel overlay (Mulas et al. 2020) (Fig. 5.A) at day 17 of the adapted protocol. This timing aligns with the established protocol for embedding CLCs in Matrigel (Fig. S2.A) and was chosen to mimic CLC maturation and morphogenesis observed in standard 3D methodologies (Sampaziotis, Brito, et al. 2015). Notably, we observed that tubes in 3D overlay cultures were significantly more prevalent (Fig. 5.D) and possessed a greater diameter when compared to standard 2D conditions (Fig. S5.A), forming extensive 3D networks. Despite the differences in morphology, expression of *KRT19* and *KRT7* remained consistent between 2D and 3D conditions (Fig. S5.B), with only early CLC marker *SOX9* exhibiting a significant change in expression. We observed decreased *SOX9* expression in 3D cultures. Given that this is an immature marker of bile duct formation, this likely indicates that there is enhanced maturation in 3D culture (Fig. S5.B). Subsequent observations showed that CLCs organised into Tubular CLCs 3 days post-overlay addition (Fig. 5.E), with tube size and branch junction frequency increasing over time. Immunostaining of day 26 Tubular CLCs further revealed co-localisation of cholangiocyte markers KRT7, KRT19 and SOX9 in tubular structures, indicative of biliary fate (Fig. 5.F). One characteristic property of *in vivo* intrahepatic bile ducts is the presence of an epithelial lumen (Sampaziotis, Brito, et al. 2015), which facilitates various secretory and absorptive processes (Brevini, Olivia C. Tysoe, and Sampaziotis 2020). We assessed this functionality using a Rhodamine-123 assay, which measures the active transport of a fluorescent substrate into the central lumen (Sampaziotis, Brito, et al. 2015; Roos et al. 2022). After 40min, Tubular CLCs demonstrated more Rhodamine-123 internalisation compared to CLCs (Fig. 5.G, Fig.S 5.D-F). We also confirmed the presence of a lumen, alongside columnar epithelial cell features such as external microvilli and intraluminal cilia, through scanning electron microscopy (Fig. 5.H). Not only did Tubular CLCs exhibit a branching morphology (Fig. 5.H), with multiple branch junctions extruding from a single tube, they also possessed buds extruding from the main tube, resembling early branch tips observed during branching morphogenesis (Lang, Conrad, and Iber 2021). Of interest, branch formation co-localised with regions of dense collagen fibrils (Fig. 5.I), suggesting a role for collagen I-binding in tube morphogenesis. To test this, we probed Tubular CLCs for collagen I-binding integrin subunit ITGβ1, observing pronounced expression around the lumen, coinciding with cholangiocyte marker expression (Fig. 5.J). Next, we investigated the effect of collagen I-specific integrin ITGα2β1 inhibition on Tubular CLC formation using small molecule inhibitor of ITGα2β1 TC-I 15 (Fig. 5.K) and ITGβ1 blocking antibody (Fig. S5.G). In the presence of TC-I-15 and anti-ITGβ1 Tubular CLCs displayed a truncated morphology, with a significant reduction in tube diameter observed in TC-I 15 conditions by day 26. Overall, our results suggest a crucial role for ITGα2β1 in bile duct morphogenesis.

## Discussion

Here, we present a protocol for the derivation of Cholangiocyte-like Cells (CLCs, Fig. 2) and Tubular Intrahepatic Cholangiocyte Organoids (Tubular ICOs, Fig. 5) from human induced Pluripotent Stem Cells (hiPSCs). This protocol leverages synthetic hydrogels with stabilised collagen I for ECM support. We also reveal the role of integrin-mediated collagen I-binding (Fig. 3) in cholangiocyte specification and intrahepatic bile duct morphogenesis. Our findings introduce a new perspective on the importance of the ECM microenvironment in cholangiocyte development within the human foetal liver. Specifically, we propose that hepatic stellate cells near the portal vein secrete high levels of collagen type I, facilitating increased fibril density and organisation to direct hepatoblast differentiation down a biliary trajectory.

Previous studies have unveiled the potential of polyacrylamide hydrogels as a substrate for hiPSC-derived hepatoblast differentiation to hepatocytes (Mittal et al. 2016) and cholangiocytes (Rizwan, Fokina, et al. 2021). However, much focus has been placed on substrate rigidity as the major inducer of differentiation, often employing standardised matrix conditions (Mittal et al. 2016; Kaylan et al. 2018). Our findings using StemBond hydrogels suggest that substrate stiffness may be secondary to stable collagen binding when determining cholangiocyte fate choice and differentiation (Fig. 2). Our revelation of the paramount importance of collagen I signalling may be attributable to the enhanced protein tethering properties of StemBond (Labouesse et al. 2021). This proposal aligns with prior findings demonstrating that ECM conjugation efficiency (S. Lee, Alice E. Stanton, et al. 2019a) and composition (Kourouklis, Kaylan, and Underhill 2016) are capable of overriding the substrate stiffness. Equally, previous studies may be limited by the difficulty to culture cells on biomimetically soft substrates. On soft substrates, previously detachment has been observed in such conditions (Mittal et al. 2016), a shortcoming that is overcome by the chemistry of StemBond. The advantageous impact of StemBond’s unique protein tethering properties is further supported by our observations of reduced hepatic fate and enhanced morphogenesis on StemBond CLCs when compared to those cultured on TCP coated with equivalent concentrations of collagen I.

Furthermore, we witnessed enhanced collagen fibril alignment on StemBond as opposed to tissue culture plastic (TCP) controls (Fig. 5). This finding is noteworthy in light of recent work by Sapudom *et al*. (2023), which places significant emphasis on the importance of ECM organisation over substrate stiffness in the induction of cellular contractility (Sapudom et al. 2023). Contractility is a major regulator of biliary differentiation (Kaylan et al. 2018; Kourouklis, Kaylan, and Underhill 2016) and has been hypothesised to be a prerequisite for YAP signalling (Rizwan, Ling, et al. 2022). Indeed, we observed an increase in YAP signalling on soft, collagen-I coated hydrogels when compared to tissue culture plastic. We also elucidated a mechanism by which hepatic marker suppression through integrin β1 binding results in enhanced biliary fate choice (Fig. 3). This is especially interesting in the context of prior research by Yang *et al*. (2017) (L. Yang et al. 2017) which demonstrates the existence of a default developmental pathway which, in the absence of transcriptional regulation, favours hepatoblast-to-hepatocyte differentiation. Building upon this, we suggest a mechanism through which the presence of dense, organised collagen I fibrils is pivotal in the inhibition of hepatic specification, enabling biliary fate choice and differentiation (Fig. 5).

The formation of consistent, large, branched networks of cholangiocytes has previously been limited to primary cells (Roos et al. 2022),(Smith et al. 2022), (Elci et al. 2024) and immortalised cell lines (Tanimizu, Miyajima, and Mostov 2007),(Chen et al. 2018). In contrast, Matrigel-embedded hiPSC-derived cholangiocytes typically yield cystic structures, with tubular formation being a less frequent occurrence (Sampaziotis, C.-P. Segeritz, and Vallier 2015),(Dianat et al. 2014),(Wu et al. 2019). Furthermore, all these studies predominantly utilise Matrigel, a substrate that lacks defined stiffness and ECM composition.

Our research stands out through, for the first time, enabling hiPSC-derived CLC tubularisation in a fully synthetic hydrogel environment. This approach offers significant advantages for biophysical studies, especially those requiring precise control over substrate stiffness and ECM composition, thus enhancing the relevance and applicability of our findings in the field. Unexpectedly, we were also able to achieve tubular structure formation on 2D StemBond hydrogels alone (Fig. 5). This finding demonstrates that 3D culture is not a prerequisite of tubularisation. This represents the first time to our knowledge that marked tubulogenesis has been achieved on 2D hydrogels without the need for complex conjugation methods enabling Notch signalling activation (Rizwan, Fokina, et al. 2021).

In conclusion, our findings mark an advance in the understanding of the importance of ECM for developmental biology and regenerative medicine. The precise control over ECM offered by our system allows for detailed exploration of the mechanical signalling impacting cholangiocyte development and bile duct morphogenesis. Thus, through providing insight into the mechanisms at play, our model paves the way for the development of innovative treatments that target biomechanical factors in diseases of the bile ducts, revolutionising the therapeutic landscape for liver disease.

## ACKNOWLEDGEMENTS

We thank Irina Moutsopoulo, Andi Munteanu and Bioinformatics Core staff for the analysis of the bulk-RNA sequencing data; Darren Clements, James Boyd and James Rands of Imaging Core for their contributions to SEM and SHG imaging; James Millar for his contribution to collagen fibril imaging; Maike Paramor, Liviu Pirvan and Vicki Murray of Genomics Core for their assistance in performing bulk RNA-seq; Floris J.M. Roos for the provision of primary cholangiocyte RNA samples; Brandon T. Wesley and Charlotte Grey-Wilson for access to the scRNA-seq analysis dataset; Tom Wyatt and Carla Mulas for agarose fibrin hydrogel optimisation; Christopher Gribben for the provision of human foetal liver sections; and Vasileios Galanakis for provision of primary hepatoblast organoids. This work was supported by the Engineering and Physical Sciences Research Council (EPSRC) Centre for Doctoral Training (CDT) in Sensor Technologies for a Healthy and Sustainable Future PhD Studentship (EP/S023046/1, to I.G.T.), Wellcome Leap Grant (ref: HOPE, to L.V.), Chan Zuckerberg Initiative (CZI) (to C.M.M. and L.V.), the European Research Council (ERC) Advanced Grant (New-Chol, 741707, to L.V.), and the European Research Council (ERC) Consolidator Grant (CellFateTech, 772798, to K.J.C.).

## Author Contributions

I.G.T., L.V. and K.J.C. designed the research. I.G.T. carried out the bulk of the experiments and data analysis. I.G.T., L.V. and K.J.C. wrote the paper. C.M.M. performed primary foetal liver tissue acquisition and hepatoblast organoid experiments. D.D. performed optimisation of hepatoblast-like cell seeding on hydrogels. L.C. performed agarose-fibrin optimisation experiments. A.H. patented the hydrogel technology. L.V. and K.J.C. acquired the funding. L.E. performed SEM imaging. All of the authors discussed the results and manuscript.

## Declaration of Interest

The authors declare no competing interests.

## Materials and Methods

### Polyacrylamide hydrogel synthesis

StemBond^*TM*^ polyacrylamide hydrogels were prepared as previously described (Labouesse et al. 2021). Briefly, prior to preparing the 2D hydrogels glass coverslips (VWR) were washed in 70% ethanol for 10 minutes and then rinsed in deionised water. Next, coverslips were treated with 0.2M NaOH for 20 minutes to facilitate the binding of hydroxyl (−OH) groups to the coverslip surface. The coverslips were dipped in MiliQ to remove salt residues and then dried using lint-free tissues (KimWipes, Midland Scientific) and placed in Petri dishes lined with hydrophobic parafilm. One side of each coverslip was coated with 5% BindSilane (PlusOneT M, GE Healthcare) dissolved in 10% acetic acid in ethanol (130µl and 85µl for 20mm and 14mm coverslips respectively) incubated at room temperature covered for 2 hours and then uncovered for half an hour to allow the BindSilane to evaporate. This enables the presentation of heterobifunctional linkers which allow polyacrylamide to covalently bind to the glass coverslips. After incubation, the coverslips were washed 3 times in absolute ethanol and wiped dry with KimWipes. StemBond hydrogel solutions of various stiffness were prepared according to Table 1 and subsequently desiccated for 2 and a half hours to prevent delamination. Polymerisation was initiated upon the addition of 0.5% N,N,NN - Tetramethylethylenediamine (TEMED, Sigma) and 0.1% Ammonium persulfate in miliQ (APS, Sigma). The polymerising hydrogel solution was pipetted onto a polymer film (intermediate polymer stamps, IPS) to enable droplet formation and coverslips were placed bind silane side down onto the droplets to produce a flat surface. Gel mixture volumes of 155µl, 80µl and 25µl were used for 32mm, 20mm and 14mm coverslips respectively to produce hydrogels of a height >180µm. This is above the threshold at which cells can sense the underlying stiffness of the glass coverslip (Buxboim et al. 2010). After 15 minutes the gels were fully polymerised, and the coverslips were removed from the IPS. The gels were then stored in PBS + penicillin/streptomycin (pen/strep) at 4^º^C until activation. The acrylamide to bis-acrylamide ratio was chosen to achieve the target stiffness and the concentration of 80mM 6-acrylamidohexanoic acid (AHA) was picked to enable maximum ECM binding.

**Table 1.** Gel mix composition for StemBond hydrogels. Volumes shown are per 500*µ*l aliquot. The 2M stock of 6-acrylamidohexanoic acid (AHA) was prepared in methanol.

| Volume (μl) | Supplier | Soft | Stiff |
| --- | --- | --- | --- |
| Acrylamide (40%) | Sigma | 34.7 | 48 |
| Bis-acrylamide (2%) | Sigma | 26.1 | 36 |
| AHA (2M) | Tokyo Chemical Industry | 20 | 20 |
| MiliQ H<sub>2</sub>O | On-site water purifier | 411.7 | 388.5 |
| TEMED | Sigma | 2.5 | 2.5 |
| APS (10%) | Sigma | 5 | 5 |

### Hydrogel functionalisation

Before cells can adhere to the hydrogel, it needs to be functionalised with the target ECM protein (Collagen I from rat tail, Sigma). This functionalisation was performed using 1-ethyl-3-(−3-dimethylaminopropyl) carbodiimide (EDAC) N-hydroxysuccinimide (NHS) cross-linker chemistry. To enable this, hydrogels were first equilibrated with MES buffer (0.1M MES hydrate, 0.5 M NaCl, pH 6.1). The buffer was then aspirated and gels were covered with activation solution (0.5M NHS, 0.2M EDAC in MES buffer) and incubated at room temperature for 30 minutes. EDAC reacts with carboxyl groups to form unstable o-acylisourea active ester intermediates. NHS then stablises these intermediates by converting them to amine-reactive NHS esters. This enables activation of the terminal carboxyl groups of AHA. The activation solution was subsequently removed and gels were washed with pre-chilled PBS and HEPES buffer (pH 8.5) before being coated with the target protein diluted in HEPES buffer to the desired concentration. Gels were then incubated at 4^º^C overnight or for 2 hours at room temperature. Primary amines on the protein displace the active intermediate by nucleophillic attack, facilitating the formation of strong covalent bonds between the activated carboxyl groups and primary amines. Afterwards, the protein was removed and blocking solution (0.5M ethanolamine in HEPES buffer) was added to the gels to block any unbound activated groups. Gels were incubated for 30 minutes at room temperature, washed with PBS and stored in PBS + pen/strep at 4^º^C until required for cell culture. Before seeding StemBond hydrogels were equilibrated in the desired media (E8 or HepatoZYME supplemented with HGF and OSM) overnight at 37^º^C.

### Synthesis of agarose-fibrin gels

Agarose-Fibrin hydrogels are synthesised using a modified version of the matrix screening protocol specified in Mulas *et al*. (2020) (Mulas et al. 2020). Briefly, 0.02g of low melt agarose (LMA) was diluted in 500*µ*l (2% agarose) of MiliQ water and then incubated at 80^º^C for a minimum of 1 hour to allow the agarose to dissolve. Once the agarose was dissolved it was transferred to 37^º^C and kept there until use. In sterile conditions the agarose gel media mix (Table 2) was combined with the 500*µ*l of dissolved agarose and the gel mix was placed in a 12 well plate. Once the gel had begun to polymerise then 40*µ*l of thrombin was pipetted on top of the gel in order to cross link the fibrinogen. The gel was then allowed to polymerise at 4^º^C for 15 minutes.

**Table 2.** Gel mix composition for collagen I agarose-fibrin hydrogels. Volumes shown are per 1ml aliquot.

| Gel |  | Media Mix Composition |  |  |  |
| --- | --- | --- | --- | --- | --- |
| Thrombrin (μl) | Agarose (μl) | Fibrinogen (μl) | Collagen I (μl) | Basal Media (μl) | Total Volume (μl) |
| 20 (1 U/ml) | 500 (2% volume) | 30 (0.15mg/ml) | 250 (1mg/ml) | 180 (18% volume) | 1000 |

### Human iPSC maintenance

The FSPS13B hiPSC cell line was maintained in Essential 8 medium supplemented with basic fibroblast growth factor (bFGF/FGF2, 25ng/ml) and TGFβ1 (2ng/ml) on 6-well plates coated with 5µg/ml Vitronectin-XF (STEMCELL Technologies). Once they reached 80% confluency the cells were passaged in clumps at a 1:20 dilution. Cells were passaged no later than every 5 days. Cells were detached from plates during passaging by incubation in 0.5mM EDTA (Thermo Fisher Scientific) at room temperature for 5 minutes. After EDTA removal the cells were washed once with medium and then blasted with medium to induce detachment. The cells were incubated at 37^º^C and 5% CO_2_.

### Hepatocyte-like cell and cholangiocyte-like cell differentiations

Hepatocyte and cholangiocyte differentiations were performed as previously described (Hannan, C. P. Segeritz, et al. 2013; Sampaziotis, Brito, et al. 2015; Sampaziotis, De Brito, et al. 2017; Olivia C Tysoe et al. 2019). Briefly, hiPSCs were seeded onto gelatin-MEF coated 12 well plates using ROCK inhibitor at a density of 50,000 cells/cm^2^ and left in E8 with bFGF and TGFβ1 for one subsequent day to enable cell growth. Gelatin-MEF coating was performed through 45 minute room temperature incubation of plates in 0.1% gelatin, and subsequent 37^º^C incubation overnight in mouse embryonic fibroblast (MEF) media. Differentiation was initiated and the plate was transferred to a hypoxic incubator (5% CO_2_, 5% O_2_, 37^º^C). Cells were differentiated into definitive endoderm using CDM-PVA supplemented with Activin-A (100ng/ml), BMP4 (10ng/ml), bFGF (20ng/ml), and LY294002 (10µM) for two days. Chiron (3µM) was added on day 1 and subsequently removed from day 2 onwards. DE cells were cultured in RPMI+ B27 supplemented with Activin-A (100ng/ml) and bFGF (20ng/ml) for one day and Activin-A (50ng/ml) for 5 further days to generate foregut endoderm as described previously (Hannan, Fordham, et al. 2013). The foregut endoderm was transferred to hepatoZYME with HGF (50ng/mL) and OSM (20ng/mL) for 4 more days to produce hepatic progenitor cells. The cells were then either continually cultured in hepatoZYME with HGF and OSM to generate hepatocytes, with media changes every 2 days. Alternatively cells were cultured in RPMI+B27 with RA (3µM), FGF-10 (50ng/mL) and Activin-A (50ng/mL) for 4 days and in common bile duct (CBDM) medium supplemented with 50ng/ml EGF (R&D Systems) and 10µM forskolin (FSK, Sigma) for 6 days to generate immature and mature cholangiocytes subsequently. The immature cholangiocyte medium was changed daily, whilst the mature cholangiocyte medium was changed every 2 days. Hepatocytes and cholangiocytes were fixed at day 26 or maintained up until day 33. Cell medium used is summarised in Table S1.

### hiPSC-derived CLC formation in Matrigel

At day 17 of cholangiocyte-like cell differentiation cells were dissociated from TCP into clumps and passaged Cell Dissociation Buffer (CDB, Gibco, Life Technologies) and re-suspended in CBDM supplemented with 50ng/ml EGF (R&D Systems) and 10µM FSK, which promotes intraluminal fluid secretion and organoid formation in resistant cell lines. The media containing the cell clumps was combined with Matrigel (BD Biosciences) at a ratio of 1:3 and 50*µ*l droplets of the gel mixture were pipetted onto a pre-warmed 24 well plate (Corning). The plate was incubated at 37^º^C for 5 minutes to allow the gel to solidify and was then flipped upside down and incubated for a further 5 minutes to prevent the cell clumps from sinking to the bottom of the gel. After incubation, the droplet was overlaid with the supplemented William’s E + medium containing ROCK Inhibitor (Y). Cells were kept in Y for two days before the media was replaced.

### Seeding hepatoblasts onto StemBond hydrogels

At day 10 of differentiation hepatoblasts were washed with PBS and then dissociated with TryplE (Gibco) for 5 minutes to produce a single cell suspension and plated onto either tissue culture plastic (TCP) or StemBond hydrogels coated with the desired ECM with ROCK inhibitor (Y) at a density of 500,000-750,000 cells per cm^2^.

### Tubular CLC formation in hydrogel sandwich

At day 17 of differentiation immature cholangiocytes were coated with 500*µ*l of the agarose fibrin hydrogel containing the ECM of choice, synthesised as described above and placed at 4^º^C for 15 minutes to enable gel polymerisation. This volume was determined through optimisation experiments to be the minimum amount required to fully coat the hydrogel and cells. Room temperature CBDM medium supplemented with EGF (50ng/mL) and FSK was then placed on top of the hydrogel sandwiches and cells were maintained in the incubator up until day 26.

### Derivation and maintenance of primary HBOs

Primary human foetal liver tissue was obtained with ethical approval (REC:96/085, REC:18/NE/0290) as previously described by Wesley *et al*. (2022) and dissociated into a single cell suspension (Wesley et al. 2022). HBOs were formed via re-suspension in basal heptoblast organoid medium (HBO-M); DMEM/F12 (ThermoFisher) supplemented with 1% HEPES (Sigma), penicillin–streptomycin and GlutaMAX (Thermofisher). Growth Factor Reduced Phenol Free Matrigel (Corning) was added to re-suspended cells at a 2:1 ratio of Matrigel to cell suspension to produce a final solution of 66% Matrigel. 20µl domes were pipetted into each well of 48-well plates (Corning) which were placed in the incubator upside down for 15min to enable the mixture to set and prevent cells from sinking to the bottom of the plate. Once set, 200µl of fresh complete HBO-M; basal HBO-M supplemented with 2% B27 (ThermoFisher), 20mM nicotinamide (Sigma), 2mM N-acetylcysteine (Sigma), 50% WNT3A conditioned medium, 10% R-spondin conditioned medium (in house tissue culture facility), 50ng/ml EGF (R&D), 5µM A83-01 (Tocris) and 10µM Y27632 (Selleckchem). Media was changed every two days and cells were passaged every 10-14 days at a split ratio 1:2–1:4. Prior to passaging, cells were re-suspended into clumps by mechanical dissociation of the Matrigel dome and subsequently incubated on ice for 5min per dome to allow the Matrigel mix to liquefy. Once incubated, the suspension was centrifuged at 400g for 5min and HBOs were seeded onto fresh plates using the protocol described above. Cell medium used is summarised in Table S1.

### Differentiation of primary HBOs to cholangiocyte progenitors

HBOs were cultured with HBO differentiation medium - complete HBO-M without A83-01 supplemented with TGFβ (2ng/ml, Biotechne) - for 8 days, applying fresh medium every 2 days.

### Production of Collagen I-Matrigel mix

Collagen I from rat tail (Sigma, 4mg/ml) was supplemented with 1.69% NaOH to enable collagen gel formation and kept on ice until use. When seeding HBOs the collagen solution was mixed at a 1:1 ratio with Growth Factor Reduced Phenol Free Matrigel (Corning), this mixture was then combined with the cell suspension at a 2:1 ratio during seeding following the protocol described above to produce a final collagen I concentration of 1.3mg/ml.

### Cytochrome P450 3A4 assay for hepatocyte functionality

Cytochrome P450 3A4 (CYP3A4) activity was measured using the P450-Glo CYP3A4 Assay (ProMega) using the reagents provided. Cells in a 12 well plate were incubated in 500µl of Luciferin reagent (diluted 1:1,000 in hepatoZYME) at 37^º^C for 1 hour in the dark. A negative control containing Luciferin, but no cells was also incubated during this period. Post-incubation the diluted Luciferin and a detection reagent were added to an opaque white 96 well plate at a 1:1 ratio (50µl of each reagent) in technical triplicates. Samples underwent a 20min equilibration at room temperature in the dark. A gloMAX plate reader was used to record luminescence, with brighter fluorescence corresponding to greater CYP3A4 activity. After the assay, cells were split using TryplE and the total cell number per sample was counted. The positive control was substracted from all samples and results were normalised per million cells measured as relative light unit per ml normalised to every million cells (RLU/ml/Million).

### Gamma-Glutamyltransferase assay for cholangiocyte functionality

Gamma-Glutamyl Transferase activity (GGT) was measured using the Colorimetric Gamma-Glutamyl Transferase Assay Kit (AbCam) following the provider’s instructions. 500*µ*l of a colorimetric substrate was added to cells in a 12 well plate and the plate was incubated for 1hr at 37^º^C until a colour change was visible. A negative control with no cells was also incubated. After incubation the colorimetric substrate was added to the transparent 96 well plate (100*µ*l/well) in technical triplicates. A standard curve of protein concentration was prepared using p-nitroaniline (pNA) dilutions of 0-40 nmol/well. The optical density (OD) of samples was measured at a wavelength of 418nm using a Molecular Devices SpectraMax M2 Multimode Plate Reader. Cells were split using TryplE and the total cell number was counted to normalise results per million cells. Readings were normalised to the 0 standard and then plotted against the standard curve to determine the amount of pNA in the samples. GGT activity (nmol/min/ml) of samples was then calculated using the following equation.

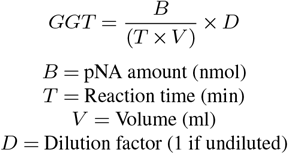

### Rhodamine-123 transport assay for cholangiocyte morphogenesis

Tubular CLC functionality was measured via transport using the Multi Drug Resistance (MDR)-1 transporter was measured by the Rhodamine-123 assay as previously described (Roos et al. 2022; Sampaziotis, Brito, et al. 2015). Briefly, cells were washed with PBS prior to 5min incubation in Rhodamine-123 (100 µM, Sigma Aldrich) at room temperature. Cells were then washed 1x using basal common bile duct medium (CBDM) and incubated (5% CO_2_, 5% O_2_, 37^º^C) for 40min to enable internalisation. Nuclei were counterstained using DAPI (3mM, Thermofisher) and cells were subsequently imaged using a confocal microscope (LSM 980 with Airyscan 2). Percentage Rhodamine-123 coverage was measured in tiled images of whole wells (2.5x magnification, 20 tiles per well) across biological replicates (*n*=3). Quantification was performed in CellProfiler. In Fiji, fluorescence measurements were made of the tube/organoid interior and exterior to measure internalisation and normalised to the maximum measurement to account for background.

### Inhibition of collagen binding

Functionality of the integrin *β*1 receptor was blocked using the Anti-Integrin *β*1, clone AIIB2 antibody (Merck) which has been previously validated to work for this purpose (Hutton et al. 2010). AIIB2 (2*µ*g/ml) was added to cells three days post-seeding (day 13 for CLCs, day 3 for HBOs) to allow time for recovery and subsequently re-applied with each media change until termination of differentiation. To control for antibody-binding anti-human IgG (AbCam) was added to cells at the same concentration. Specific binding of cells to collagen I using integrin *α*2*β*1 was blocked using the small molecule inhibitor thiazolidine-modified compound 15 (TC-I-15, Tocris Biosciences) (Hunter et al. 2021) at a working concentration of 50µM repeating the protocol discussed above. Tube diameter measured in Fiji was used as a metric of Tubular CLC formation and quantified across biological replicates (*n*=3).

### Immunofluorescence (IF)

Cells were fixed in 4% paraformaldehyde (PFA) for 15 minutes, washed twice in PBS and stored at 4^º^C until further use. When required for immunostaining cells were permeabilised with 0.1% Triton X-100 in PBS for 15 minutes, washed twice with PBS, and subsequently blocked in 10% Donkey Serum (DS) for 2 hours, both incubations were performed at room temperature. Primary antibodies diluted in 10% DS at the dilution recommended by the supplier (complete list provided in Table S2) were added to the cells and incubated overnight at 4^º^C on the rocker. The next day cells were washed four times in PBS and subsequently incubated in secondary antibodies diluted in 10% DS to the concentration recommended by the supplier (1:1000). Incubation occurred over 2 hours at room temperature. After incubation cells were washed twice in PBS and the nuclei were stained with DAPI (1:10,000, diluted in PBS) for 15 minutes at room temperature. Post-staining, cells were washed three times in PBS and stored in PBS at 4^º^C until required for imaging. Gels were mounted in 35 mm µ-Dishes (IBIDI) before imaging. Mounting was performed through pipetting 10ul PBS into the centre of the dish and subsequently inverting the gel onto the dish.

### Immunohistochemistry of paraffin embedded samples

Human foetal tissue was acquired from patients undergoing elective terminations (ethical approval from East of England—Cambridge Central Research Ethics Committee REC-96/085) as previously described (Wesley et al. 2022), embedded in paraffin and sliced into 4*µ*m thick sections. Previously fixed and paraffin embedded sections were obtained from Carola Maria Morell of the Vallier lab. Slides were washed 2x in Histoclear II histology clearing agent (SLS, NAT1334) for 10 minutes to remove paraffin penetrated into the tissue. Deparaffinated slides were then washed 3x for 5 minutes each in 100% ethanol to rehydrate them and 1x in water for 5 minutes. Slides were then steamed for 20 minutes in the buffer of choice (1x Citrate buffer pH 6, Sigma; 1mM EDTA pH 8, ThermoFisher) to enable antigen retrieval and left at room temperature for another 20 minutes. Samples were washed with water and then PBS 0.1% TritonX-100 (Sigma, T8787) for 5 minutes each. Prior to staining, samples were blocked with PBS 10% donkey serum (DS, Biorad, C06SB) 0.1% TritonX-100 at room temperature for 1hr. The blocking solution was removed by tapping the slide gently on one side, and the primary antibody of choice in PBS 10% DS 0.1% TritonX-100 (diluted to the desired concentration) was left on the slides overnight at 4^º^C in a humid environment to prevent evaporation. The next day the primary antibody was removed and slides were washed 3x for 5 minutes each in PBS 0.1% TritonX-100 before incubation in the desired secondary antibody in PBS 10% DS 0.1% TritonX-100 (1:1,000) at room temperature for 1hr. Afterwards, samples were washed 2x for 5 minutes in PBS 0.1% TritonX-100 and 1x in PBS 0.1% TritonX-100 Hoechst (1:10,000, Sigma B2883-100mg) to co-stain the nuclei. Hoescht was removed by washing the samples 2x in PBS for 5 minutes. Slides were dried and then developed by the addition of Fluoromount-G Mounting Medium (Invitrogen) and a top coverslip which was sealed using clear nail polish.

### Quantitative reverse transcription polymerase chain reaction (RT-qPCR)

Once the cells reached the desired stage of differentiation, they were lysed in the well using RLT buffer (Qiagen, 74134). The extracted RNA was converted to cDNA by SuperScript IV Reverse Transcriptase (ThermoScientific, 18090010). For RT-qPCR, pre-designed primers (complete list provided in Table S3) were combined with the SybrGreen mastermix (ThermoScientific, NC0903497) and RT-qPCR was performed on the Applied Biosystem Step One machine. Endogenous expression was assessed using the reference genes GAPDH, PBGD and RPLP0.

### Flow cytometry

Cells were incubated at 37^º^C in TryplE (Gibco) for 5 minute intervals followed by pipetting in order to generate a single cell suspension. Cell suspensions were filtered to remove clumps using a 40um cell strainer (Starlab). They were then washed with basal hepatoZYME and placed in the fixative 4% paraformaldehyde (PFA, Santa Cruz Biotechnology sc-281692) for 15 minutes at 4^º^C. Post-fixing, cells were washed with 1% Bovine Serum Albumin (BSA, Merck) in PBS and re-suspended in PBS containing 1% BSA/0.2% Saponin (Merck) to permeablise and block at room temperature for 30 minutes or overnight at 4^º^C. Primary antibodies were added to the PBS (1% BSA, 0.2% Saponin) at a 1:100 dilution and cells were incubated overnight at 4^º^C. Post-incubation, the cells were re-suspended in fresh PBS (1% BSA, 0.2% Saponin) and secondary antibodies were applied at a 1:400 dilution. The cells were incubated in the secondary antibody for 1 hour at room temperature in the dark. Finally, cells were washed with PBS (0.1% BSA), re-suspended in PBS (0.1% BSA) and transferred to a FACS tube (Corning). Flow cytometry was performed using a BD LSRFortessa Cell Analyzer and a minimum of 20,000 events were recorded. FlowJo software was used to analyse flow cytometry results.

### Confocal light microscopy

Imaging of fixed samples was performed using confocal microscopes (LSM 980 with Airyscan 2 and Zeiss LSM 710), taking Z-stacks with a 8µm, 4µm or 2µm step using either a 10x, 20x or 40x objective respectively. Single stack imaging was also carried out on the Leica 6000 fluorescent microscope using a 20x objective. Image analysis was executed in Fiji (Fiji is just ImageJ) (Schindelin et al. 2012). Quantification was performed on maximum intensity projections of the Z-stacks. Quantification of the nuclear to cytoplasmic ratio was performed using a CellProfiler (Carpenter et al. 2006) pipeline by selecting nuclei in the desired channel and cells another channel within a set size threshold and subtracting the nuclei from the cell to create a mask of the cytoplasm. The mean intensity of the cytoplasm and nuclei were measured, and their ratio was calculated. This protocol was later improved using an image analysis pipeline (https://github.com/CMulas/KTR_analysis) to segment whole cells using ZO1 staining in CellPose and quantify the nuclear to cytoplasmic ratio of active YAP protein using the CellProfiler pipeline.

### Scanning electron microscopy

Scanning electron microscopy (SEM) of the hydrogels was performed in collaboration the core imaging facility using a TESCAN Clara Volume scanning electron microscope (Cambridge Stem Cell Institute imaging facility). Samples were fixed with EM grade 4% PFA and 2.5% glutaraldehyde in sodium cacodylate buffer (pH 6.5, provided by the EM core) for one hour at room temperature and overnight at 4^º^C. The fixed gels were washed 3x with ddH20 for 10 minutes each. Washed samples were then dehydrated using a graded ethanol series (30%, 50%, 70%, 90% and 100%) for 10 minutes each before an additional 2x 100% ethanol washes. Samples were then dried in a critical point drier which prevents sample shrinkage and preserves delicate surface structures by avoiding tangential forces. The samples were next coated with 8nm of gold sputter to allow for dissipation of electrons from the SEM. The more electrons the sample can dissipate the higher voltage and beam current can be used – allowing for better resolution in the SEM. Samples were mounted onto a sticky double sided carbon tab and an aluminium SEM stump subsequently and imaged using the Everhart-Thornley detector (E-T detector) which images secondary electrons excited from the sample by the electron beam. SEM images were false coloured using Adobe Photoshop (version 25.1.0).

### Second harmonic generation

Second harmonic generation (SHG) of collagen in histological paraffin-embedded human foetal liver sections was performed with assistance from the core imaging facility using the Zeiss LSM 880 with Non-Linear Optics (NLO) inverted confocal microscope (Cambridge Stem Cell Institute imaging facility) using a water immersion 20x objective lens (W Plan-Apochromat 20x/1.0 DIC VIS-IR objective). Collagen fibrils were detectable between emission wavelengths of 450-480nm using an ultrafast pulse Spectra-Physics Insight laser. Prior to imaging paraffin was removed and samples were re-hydrated as discussed above. SHG signal was differentiated from autofluorescence and background noise, defined as a broad spectra, through unmixing in Zeiss ZEN Black 2.3 SP1 software.

### Bulk RNA-sequencing

RNA samples of cholangiocyte-like cells differentiated in soft/stiff collagen I StemBond and CLC control conditions were extracted as described above and submitted to the in-house genomics core facility for RNA quality control and fragmentation to check for partial degredation and produce short sequences (40-400bp) capable of being read by the sequencing platform. Preparation of the cDNA library was performed by the in-house facility using Pico mammalian V2 (Takara, USA), NuGen (NuGen, CA, USA) or RiboZero and Nextflex (Bioo Scientific, TX, USA) kits. Bulk sequencing was executed on Illumina HiSeq4000.

### Bioinformatics

Analysis of bulk-RNA transcriptomic datasets was performed by the Cambridge Stem Cell Institute core bioinformatics facility. Genes of interest were queried and plots were produced using an online Shiny application. GSEA output of the Shiny application is summarised in Tables S4 and S5.

### Analysis of pre-generated scRNA-sequencing data

A sc-RNA transcriptomic dataset of cholangiocyte-like-cell differentiation *in vitro* was obtained from Brandon Wesley (Wesley et al. 2022). Analysis was carried out using the ScanPy (single cell analysis in Python) Python toolkit (https://scanpy.readthedocs.io/en/stable/generated/scanpy.pl.pca.html) (Wolf, Angerer, and Theis 2018) following the protocol descriped by Wesley *et al*. (2022).). Briefly, quality control was performed by excluding cells expressing fewer than 2500 genes or more than 0.1% mitochondrial content. The data matrix was then normalised to 10,000 reads per cell so that counts become comparable among cells, logarithmised and highly variable genes were identified. Then the effects of total counts per cell and the percentage of mitochondrial genes expressed were regressed out, the data was scaled to unit variance and values exceeding a standard deviation of 10 were clipped. Calculation of differentially expressed genes was performed via T-test using a *P*-value threshold (*P*< 0.01) and absolute log-fold change threshold (log-fold change> 1) as specified in Wesley *et al*. (2022) (Wesley et al. 2022) and genes were filtered for differential expression. Once filtered, the genes of interest were queried against the dataset to produce PCA plots and heatmap representations. Plots and heatmap representations of datasets were generated using Matplotlib.

### Statistical analyses

Statistical analysis was performed using GraphPad Prizm (version 10.0.2) or RStudio (version 2023.09.1+494). For comparison of two mean values a 2-Sided Student’s T-test or a Mann-Whitney test was used as a non-parametric equivalent when values did to conform to normality or equal variance assumptions. For comparison of multiple values either a One-Way ANOVA/non-parametric Kruskal-Wallis test was used with Tukey’s or Dunn’s multiple comparisons test respectively to correct for multiple comparisons or multiple T-tests were used. When comparison to a standard set value was required for multiple samples (e.g. when measuring fold change expression in multiple markers) a One Sample T-test or non-parametric Wilcoxon Signed-Rank test was used. Biological replicates are specified in figure legends and statistical analysis was performed when biological replicates were ≥ 3. A *P*-value of ≤ 0.05 was considered significant as determined by the tests above and outlying data-points were identified using Grubbs test. To smooth the curves generated in figures 3.B and 5.G LOWLESS curve fitting was performed in GraphPad Prizm using 10 points in the smoothing window. Graphs in RStudio were generated using plugins ggplot2 (https://ggplot2.tidyverse.org/) and ggpubr (https://rpkgs.datanovia.com/ggpubr/).

To generate coherency measurements of collagen fibril organisation the Fiji plugin OrientationJ (http://bigwww.epfl.ch/demo/orientationj/) was used to quantify fibril orientation and coherency. Differentiated HBO outgrowth was measured using the NeuronJ (https://imagescience.org/meijering/software/neuronj/) plugin to trace branches and subsequently quantify branch length. Gene set enrichment analysis (GSEA) of gene lists obtained through the bioinformatics facility Shiny application (https://github.com/Core-Bioinformatics/bulkAnalyseR) and plotting of GSEA results was performed via R plugins fgsea (https://github.com/ctlab/fgsea), msigdbr (https://igordot.github.io/msigdbr/), dplyr (https://dplyr.tidyverse.org/), org.Hs.eg.db (https://bioconductor.org/packages/release/data/annotation/html/org.Hs.eg.db.html), clusterProfiler (https://guangchuangyu.github.io/software/clusterProfiler/) and enrichplot (https://github.com/YuLab-SMU/enrichplot).

## Abbreviations

α-ITGβ1: Integrin Beta 1 Blocking Antibody
AFP: Alpha-fetoprotein
AHA: 6-acrylamidohexanoic acid
ALB: Albumin
CLCs: Cholangiocyte-Like Cells
COL I: Collagen Type I
COL IV: Collagen Type IV
CYP3A4: Cytochrome P450 3A4
DP: Ductal Plate
ECM: Extracellular Matrix
FCM: Flow Cytometry
GGT: Gamma-Glutamyl Transferase
GSEA: Gene Set Enrichment Analysis
HA: Hepatic Artery
HBO-M: Hepatoblast Organoid Medium
HBOs: Hepatoblast Organoids
HFL: Human Fetal Liver
hiPSC: Human Induced Pluripotent Stem Cells
HLCs: Hepatocyte-Like Cells
HepSCs: Hepatic Stellate Cells
ICOs: Intrahepatic Cholangiocyte Organoids
IF: Immunofluorescence
IHC-P: Immunohistochemistry-Paraffin
IHBDs: Intrahepatic Bile Ducts
ITGα2: Integrin Alpha 2
ITGαV: Integrin Alpha V
ITGβ1: Integrin Beta 1
KRT19: Keratin 19
KRT7: Keratin 7
MMP: Matrix Metalloproteinase
PCW: Post-Conception Weeks
PCA: Principal Component Analysis
PPARA: Peroxisome Proliferator-Activated Receptor Alpha
PPARD: Peroxisome Proliferator-Activated Receptor Delta
PPARG: Peroxisome Proliferator-Activated Receptor Gamma
PV: Portal Vein
RT-qPCR: Real-Time Quantitative Polymerase Chain Reaction
SEM: Scanning Electron Microscopy
SERPINA1 / A1AT: Alpha-1 Antitrypsin
SHG: Second Harmonic Generation
SOX9: SRY-Box Transcription Factor 9
TAZ: Transcriptional Coactivator with PDZ-Binding Motif
TC-I-15: Thiazolidine-Modified Compound 15
TCP: Tissue Culture Plastic
TGFβ: Transforming Growth Factor Beta
VIM: Vimentin
YAP: Yes-Associated Protein

## Supplementary Information

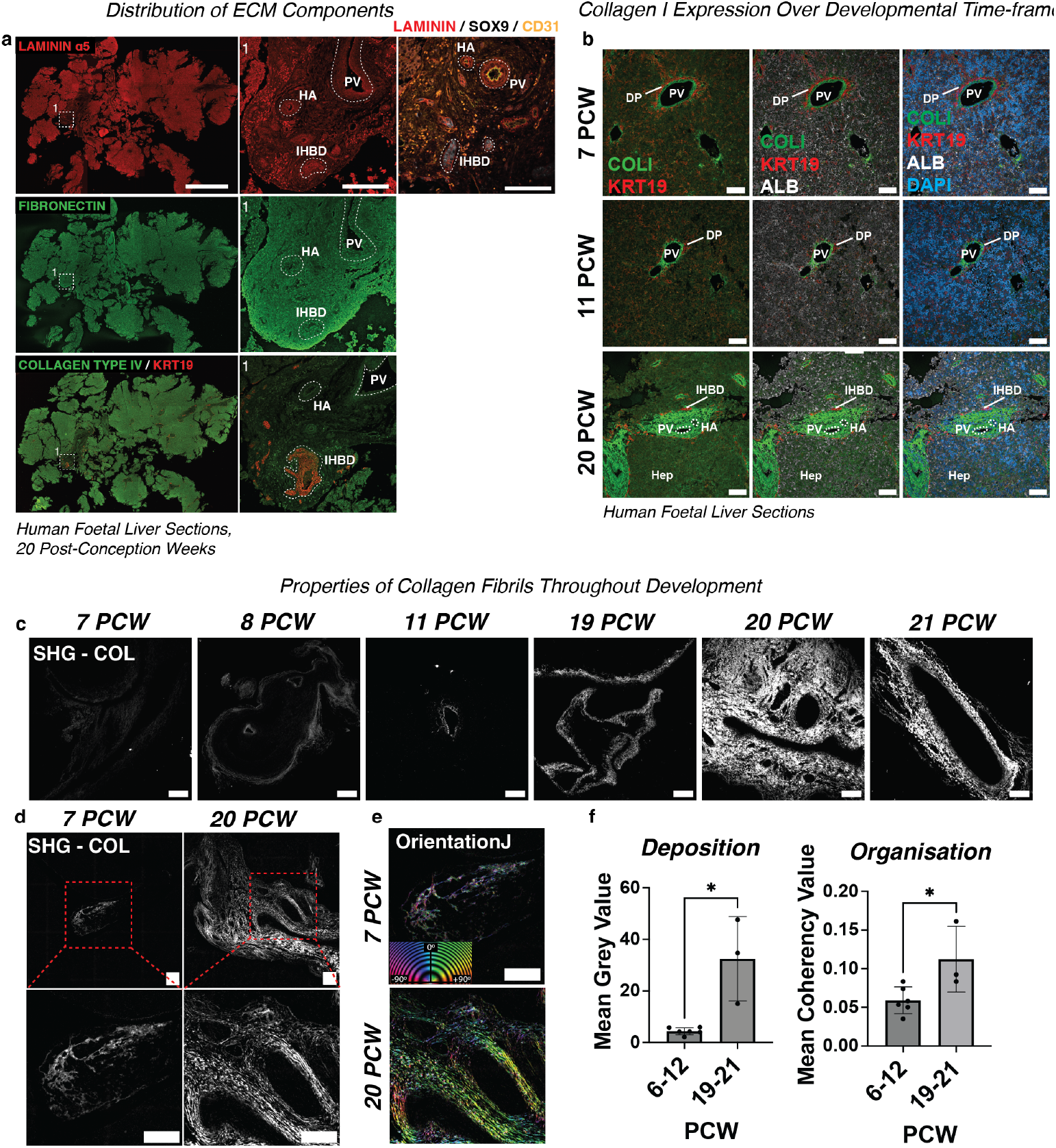
Collagen type I is deposited at the portal vein throughout *in vivo* human liver development. **(A)** Immunostaining of entire human foetal liver section at 20 post-conception weeks (PCW) for fibronectin, laminin and collagen IV alongside biliary markers SOX9 and KRT19 and vessel marker CD31. Scale bars: 2mm and 250µm, respectively. Both staining for laminin *α*5 and overall laminin content (denoted laminin) are shown. HA: hepatic artery, PV: portal vein, IHBD: intrahepatic bile duct. **(B)** Immunostaining of human foetal liver sections throughout human liver development. Stains of collagen I (COL I, green), cholangiocyte marker Keratin 19 (KRT19, red) and hepatocyte marker Albumin (ALB, green) at 7, 11 and 20 PCW shown with and without nuclear stain DAPI (blue). Scale bar: 100µm. DP: ductal plate. **(C-F)** Second Harmonic Generation (SHG) visualisation of collagen fibrils over developmental timeframe (in PCW). Scale bars: 500µm. **(C)** Sections at 7, 8, 11, 19, 20 and 21 PCW. **(D)** 7 and 20 PCW sections shown. Region of interest indicated in red. **(E)** Visualisation of fibril orientation using the OrientationJ Fiji Plugin. Colour indicates degree of orientation (green = more aligned) and intensity coherency value. Scale is 500µm. **(F)** Quantification of fibril deposition (intensity measured as mean grey value) and organisation (coherency value) over developmental timeframe (measured in PCW) across biological replicates (6-12: *n*=5, 19-21:*n*=3), Mann-Whitney test). Asterisk indicates *P*-value: *≤0.05.

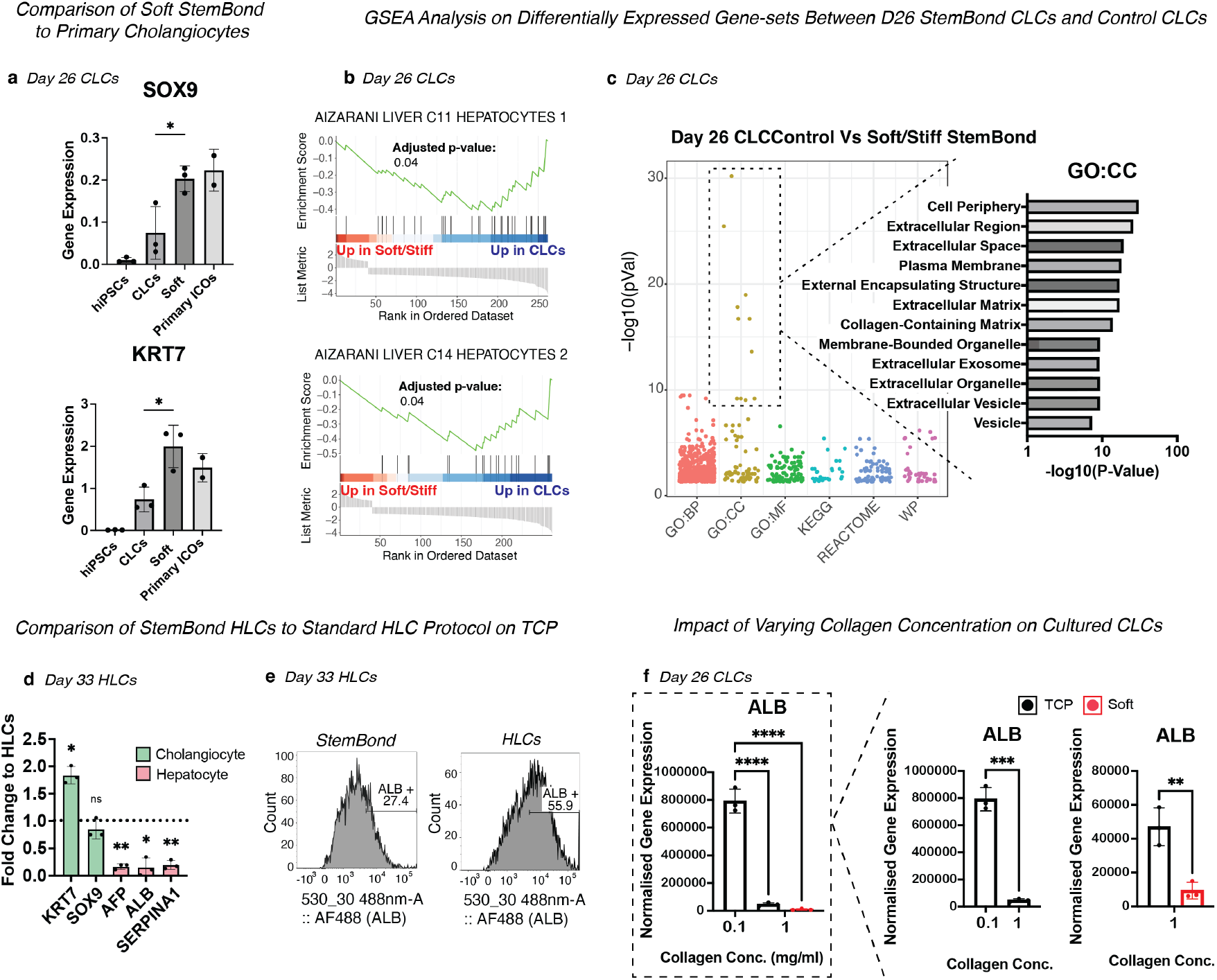
Collagen I-rich StemBond hydrogels direct hiPSC-derived hepatoblast-like cells towards a biliary fate (A-C) and away from a hepatic fate (D-K) *in vitro*. **(A)** RT-qPCR showing mean gene expression of *KRT7* and *SOX9* in day 26 cholangiocyte-like cells (CLCs) cultured using our adapted StemBond protocol (*n*=3) or the CLC protocol (*n*=3) described by Sampaziotis *et al*. (2017) (Sampaziotis, De Brito, et al. 2017) and primary intrahepatic cholangiocyte organoids (*n*=2). Gene Expression (One-Way ANOVA) is shown relative to housekeeping genes (*PBGD, RPLP0, GAPDH*). Asterisk indicates *P*-value: *≤0.05. Error bars denote standard deviation. Primary ICOs: primary intrahepatic cholangiocyte organoids. **(B)** Gene set enrichment analysis (GSEA) of differentially expressed genes (log2FC≥1, Adjusted *P*-value ≤0.05) between day 26 cells in soft (*n*=3) / stiff (*n*=2) StemBond and control CLC conditions (*n*=2). Normalised enrichment score and *P*-value are shown. Gene sets represent target cell type. **(C)** Plot showing GSEA of genes that significantly differ (log2FC≥1, Adjusted *P*-value ≤0.05) between control CLCs and CLCs cultured on soft and stiff StemBond hydrogels (*n*=2/4) using the Gene Ontology: Cellular Compartment (GO:CC) database. -log10(*P*-values) are shown. **(D)** RT-qPCR showing mean fold change gene expression of hepatic markers (*ALB, AFP* and *SERPINA1*) and biliary markers (*KRT7, SOX9*) in day 33 hepatocyte-like cells (HLCs) (*n*=3, One-Sample T-test) relative to housekeeping genes (*PBGD, RPLP0, GAPDH*). Asterisk indicates *P*-value: *≤0.05, **≤0.01. Error bars denote standard deviation. HLCs: gelatin TCP protocol. **(E)** Flow cytometry analysis of ALB-positive cells across StemBond and HLC conditions. Value indicates percentage of positive cells. **(F)** RT-qPCR showing mean gene expression of ALB in day 26 CLCs (*n*=3, Unpaired T-test) relative to housekeeping genes (*PBGD, RPLP0, GAPDH*) and normalised to the human induced pluripotent stem cell (hiPSC) control in CLCs cultured on different substrates. Black bars represent collagen TCP and red bars soft StemBond. Asterisks indicate *P*-value: **≤0.01, ***≤0.001, ****≤0.0001. Error bars denote standard deviation. TCP: tissue culture plastic.

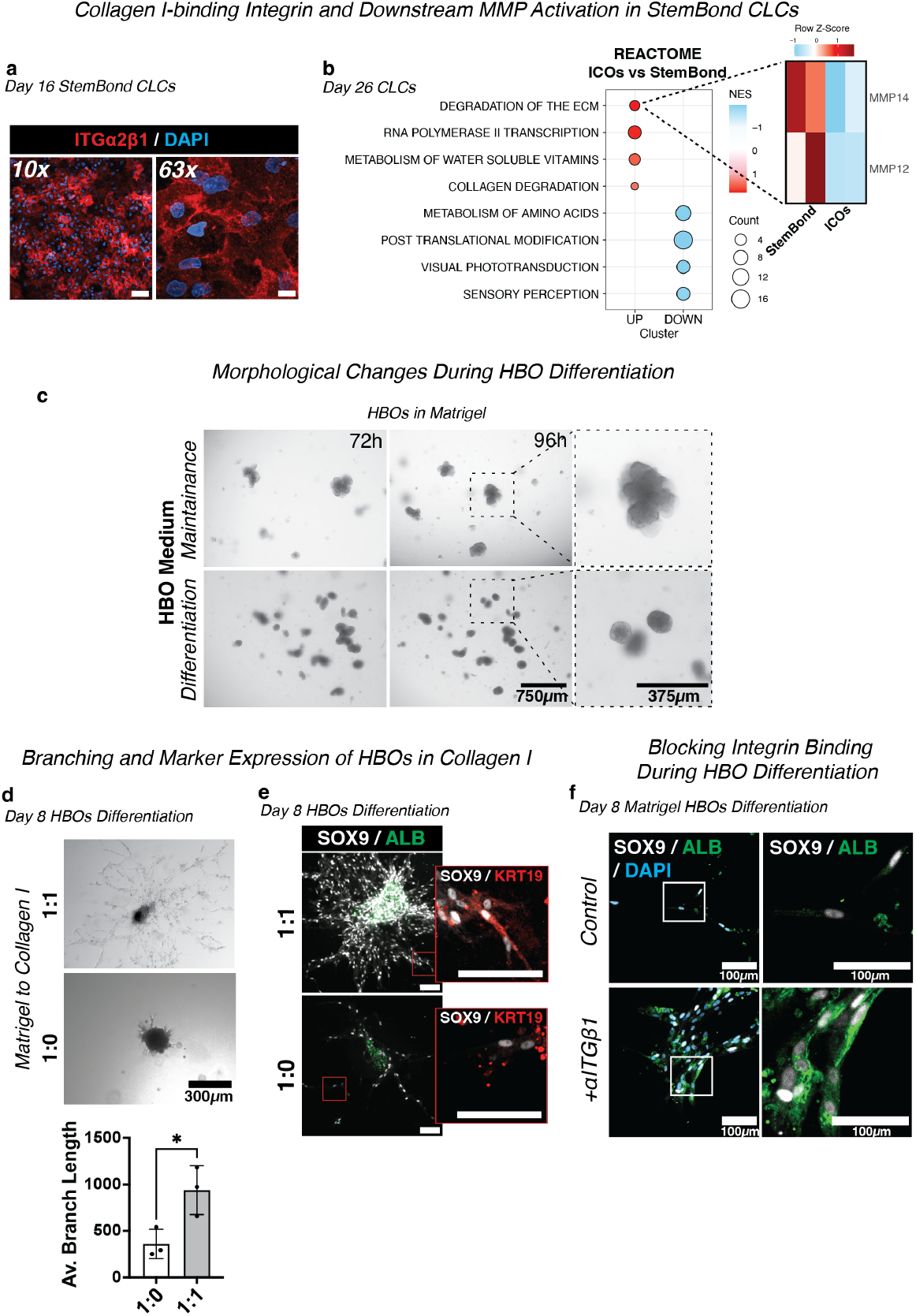
Collagen type I-binding suppresses hepatic differentiation in hiPSC-derived cholangiocyte-like cells (A-C) and primary hepatoblast organoids (D-G). **(A)** Immunostaining of day 16 cholangiocyte-like cells (CLCs) for integrin α2β1 (ITGα2β1, red) nuclei are co-stained with DAPI (blue). **(B)** Gene set enrichment analysis (GSEA) of differentially expressed genes (log2FC≥1, Adjusted *P*-value ≤0.05) between day 26 cells in StemBond and control CLC conditions. Heatmap represents row Z-score of genes from the top differentially expressed gene sets. Dot size indicates gene-count and colour represents normalised enrichment score. **(C)** Brightfield images of primary hepatoblast organoids (HBOs) cultured under either differentiation or maintenance conditions at 24h timepoints. Scale bars: 750µm and 375µm, respectively. **(D)** Brightfield images of day 8 differentiated HBOs cultured in a 1:1 mix of collagen I and Matrigel or control 1:0 (Matrigel) conditions. Scale bar: 300µm. Quantification of the mean branch length (in pixels) across biological replicates (*n*=3, Unpaired T-test) as measured by NeuronJ Fiji plugin is shown. Asterisk indicates *P*-value: *≤0.05. Error bars denote standard deviation. **(E)** Immunostaining of SOX9 (greyscale), ALB (green) and KRT19 (red) in day 8 differentiated HBOs cultured in a 1:0 or 1:1 mix of Matrigel and collagen I. Scale bar: 100µm. **(G)** Immunofluorescence images of day 8 differentiated primary hepatoblast organoids (HBOs) in Matrigel cultured with or without an integrin β1 blocking antibody (+αITGβ1). Cells are stained for SOX9 (greyscale), ALB (green) and KRT19 (red). Nuclei are co-stained with DAPI (blue). Scale bars: 100µm for BF and immunostaining images, respectively.

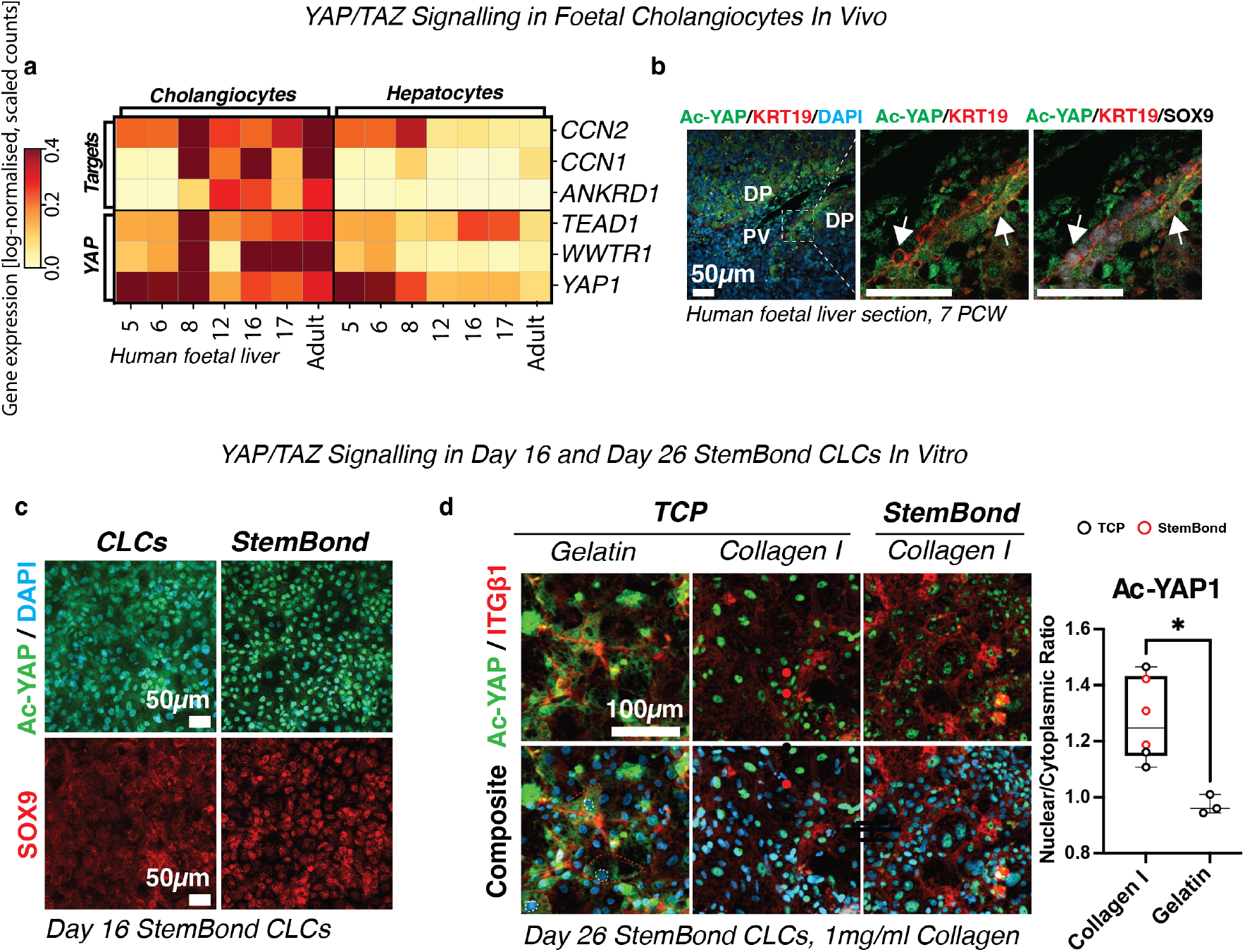
YAP/TAZ signalling is involved in *in vitro* and *in vivo* cholangiocyte differentiation. **(A)** Heatmap visualisation of all integrated single-cell transcriptomic data of foetal and adult human hepatic cells (human foetal liver cells, HFLCs) obtained via Wesley *et al*. (2022) (Wesley et al. 2022) generated using the 10x Genomics workflow; annotation indicates cell-specific lineages (top), target gene sets (left) and age in post-conception weeks (bottom). Colour indicates mean gene expression values. **(B)** Human foetal liver section at 7 post-conception weeks (PCW) stained for active YAP (Ac-YAP, green), KRT19 (red), SOX9 (greyscale) and co-stained for DAPI (blue). Arrows point to regions of co-expression. DP: ductal plate, PV: portal vein. Scale, 50µm **(C)** Immunostaining of day 16 CLCs for active YAP (Ac-YAP, green), SOX9 (red) and DAPI (blue). Scale bars: 50µm and 25µm. **(D)** Immunostaining of day 26 CLCs for active YAP (Ac-YAP, green), integrin β1 (ITGβ1, red) and DAPI (blue). Scale bar: 100µm and 25µm. Red dashed line indicates cytoplasm and white line nucleus. Quantification of mean Ac-YAP nuclear/cytoplasmic ratio was performed across biological replicates (*n*=3, Two-Way ANOVA). Asterisk indicates *P*-value: **≤0.01. Error bars denote standard deviation.

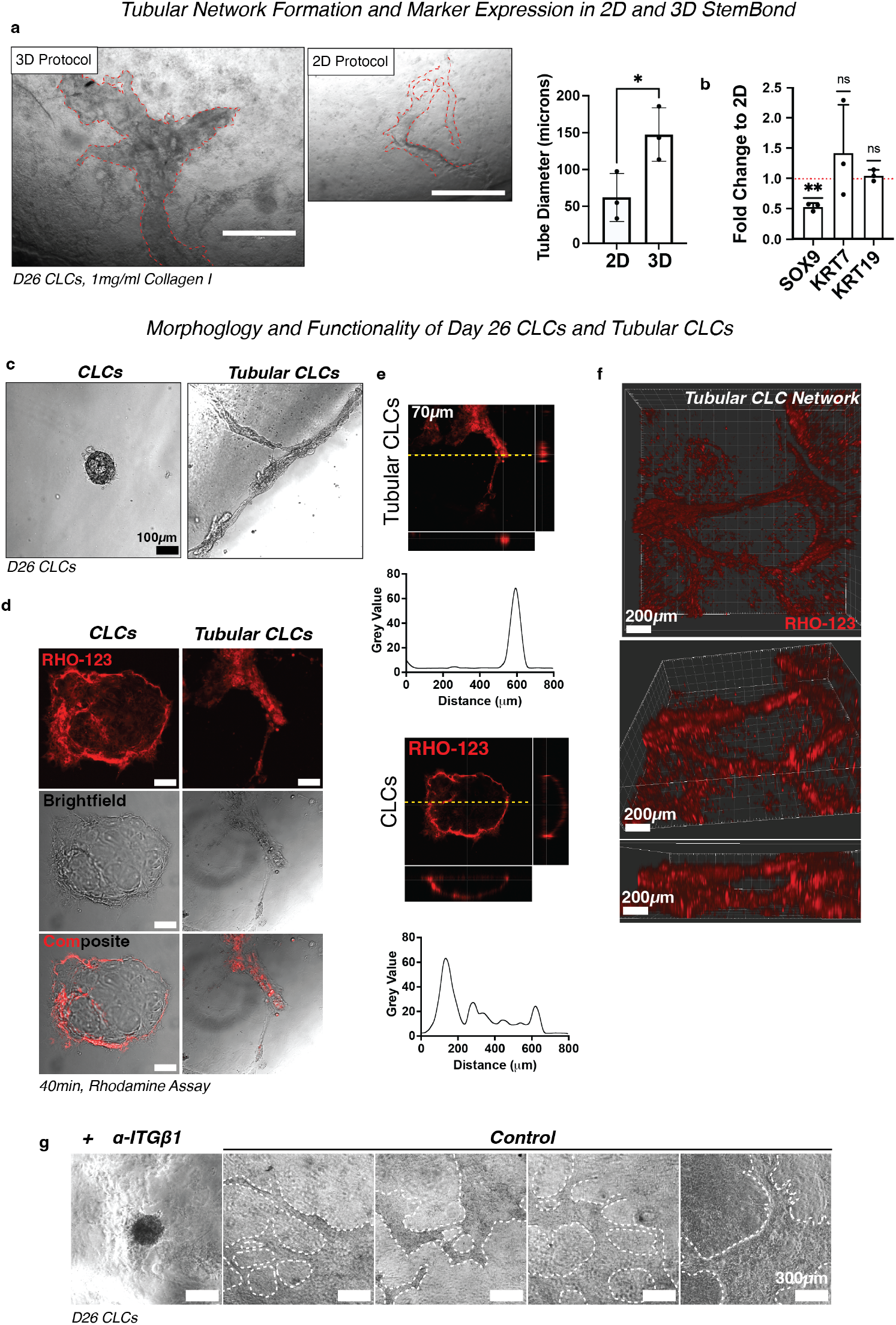
Tubular cholangiocyte-like cells are formed in StemBond-Fibrin sandwich hydrogels functionalised with collagen type I. **(A)** To scale brightfield images of day 26 Tubular cholangiocyte like cells (Tubular CLCs, outlined) in 2D and 3D StemBond conditions. Scale, 750µm. Graph represents quantification of tube diameter (mean ± SD) across biological replicates (*n*=3, Unpaired T-test). Asterisk indicates *P*-value: *≤0.05. **(B)** RT-qPCR of mean cholangiocyte marker gene (*SOX9, KRT7, KRT19*) expression in day 26 Tubular CLCs (*n*=3) displayed as fold change to CLCs cultured using the 2D StemBond protocol. Values are normalised to housekeeping genes (*PBGD, RPLP0*, GAPDH). Asterisk indicates *P*-value: **≤0.01, ns≥0.05. Error bars denote standard deviation. Dashed line represents control 2D values. **(C)** Brightfield images comparing day 26 Tubular CLCs and Matrigel-embedded CLC controls (CLCs). Scale bar: 100µm. **(D)** Max projection of confocal images of Tubular CLCs and CLC controls assayed for luminal Rhodamine-123 (RHO-123) internalisation (red) after 40min. Scale bar: 80µm. Brightfield slices also shown. **(E)** Dashed line represents region of quantification. Fluorescence intensity measurements (grey value, normalised to the highest value) across the dashed line are plotted. Scale bar: 70µm. **(F)** Confocal 3D projection of Tubular CLC branching networks, visualised using RHO-123, as viewed from above, 3D and orthogonal views. Scale, 200µm **(G)** Brightfield images of day 26 Tubular CLCs (outlined) in unsupplemented control and integrin β1 blocking antibody supplemented (+ αITGβ1) conditions. Scale, 300µm.

Cell culture medium compositions. Volumes are given per 500ml of medium.

**Table S 1.** Cell culture medium compositions. Volumes are given per 500ml of medium.
| Medium | Components | Concentration | Volume | Source |
| --- | --- | --- | --- | --- |
| E6 | DMEM/F12, HEPES | 95.3% | 500ml | Gibco |
|  | Insulin-Transferrin-Selenium, 100x | 2% (2x) | 10ml | Gibco |
|  | Sodium Bicarbonate, 7.5g/L | 0.7% (51.6mg/L) | 3.6ml | Gibco |
|  | Penicillin/streptomycin, 10,000U/mL | 1% (100U/ml) | 5ml | Gibco |
|  | L-Ascorbic acid 2-Phosphate, 100x | 1% (1x) | 5ml | Sigma |
| CDM<br>PVA | Ham's F12 Nutrient Mixture | 49% | 250ml | Gibco |
|  | Iscove's Modified Dulbecco's Media | 49% | 250ml | Gibco |
|  | Chemically defined lipid concentrate, 100x | 1% (1x) | 5ml | Gibco |
|  | Penicillin/streptomycin, 10,000U/mL | 1% (100U/ml) | 5ml | Thermo |
| | 1-thioglycerol, 1.25g/mL | 0.004% ($5 \times 10^{-5}$ g/ml) | 20μl | Sigma |
| | Insulin solution human, 10mg/ml | 0.07% (7μg/ml) | 350μl | Roche |
| | Human Apo-Transferrin, 30mg/ml | 0.05% (15μg/ml) | 250μl | Roche |
|  | Polyvinyl alcohol | 1mg/ml | 0.5g | Roche |
| RPMI+ | RPMI 1640 +Glutamax | 96% | 480ml | Gibco |
|  | B27 plus insulin supplement, 50x | 2% (1x) | 10ml | Gibco |
|  | Penicillin/streptomycin, 10,000U/mL | 1% (100U/ml) | 5ml | Thermo |
|  | MEM non-essential amino acids, 100x | 1% (1x) | 5ml | Gibco |
| Hepato<br>ZYME+ | HepatoZYME-SFM | 94% | 470ml | Gibco |
|  | Chemically defined lipid concentrate, 100x | 2% (2x) | 10ml | Gibco |
|  | MEM non-essential amino acids, 100x | 2% (2x) | 10ml | Gibco |
|  | Penicillin/streptomycin, 10,000U/mL | 1% (100U/ml) | 5ml | Thermo |
|  | L-glutamine, 200mM | 1% (2mM) | 5ml | Thermo |
| | Insulin solution human, 10mg/ml | 0.14% (14μg/ml) | 700μl | Roche |
| | Human Apo-Transferrin, 30mg/ml | 0.01% (30μg/ml) | 500μl | Roche |
| CBDM+ | William's E media | 96% | 480ml | Gibco |
|  | Nicotinamide, 0.4M | 2.5% (10mM) | 12.5ml | Sigma |
|  | HEPES buffer, 1M | 2% (20mM) | 10ml | Sigma |
|  | Sodium bicarbonate (NaHCO<sub>3</sub>), 1M | 1.7% (17mM) | 8.5ml | Sigma |
|  | Sodium pyruvate (C<sub>3</sub>H<sub>3</sub>NaO<sub>3</sub>), 100mM | 0.063% (6.3mM) | 3.15ml | Invitrogen |
|  | L-ascorbic acid 2-phosphate (AA2P), 100mM | 0.2% (0.2mM) | 1ml | Sigma |
|  | Glucose, 1M | 1.4% (14mM) | 7ml | Thermo |
|  | Insulin-Transferrin-Selenium (ITS), 100x | 1% (1x) | 5ml | SLS |
| | Dexamethasone, 10mM | 1% (0.1μM) | 5ml | R&D |
|  | Penicillin/streptomycin, 10,000U/mL | 1% (100U/ml) | 5ml | Thermo |
|  | L-glutamine, 200mM | 1.06% (2.1mM) | 5.3ml | Thermo |
| MEF | Advanced DMEM/F12 | 88.1% | 450ml | Gibco |
|  | Foetal Bovine Serum (FBS), 100x | 9.8% (10x) | 50ml | PanBiotech |
|  | L-Glutamine, 200mM | 1% (2mM) | 5ml | Thermo |
|  | Penicillin/streptomycin, 10,000U/mL | 1% (100U/ml) | 5ml | Thermo |
| | 2-Mercaptoethanol, 50mM | 0.1% (50μM) | 0.5ml | Invitrogen |
| HBO | DMEM/F12 | 97% | 475ml | Gibco |
|  | Penicillin/streptomycin, 10,000U/mL | 1% (100U/ml) | 5ml | Thermo |
|  | GlutaMAX | 1% (100U/ml) | 5ml | Gibco |
|  | GlutaMAX | 1% (100U/ml) | 5ml | Gibco |
|  | HEPES buffer, 1M | 1% (10mM) | 5ml | Sigma |

List of antibodies used for immunostaining (IF), flow cytometry analysis (FCM) and immunohistochemistry (IHC-P). KRT: Cytokeratin, Rb: rabbit, Gt: goat, Ms: mouse, Dk: donkey, Rt: Rat, Mc: Monoclonal, Pc: Polyclonal, 1^º^: primary binding, 2^º^: secondary binding, Ab: antibody, LSECs: liver sinusoidal endothelial cells, ECM: extracellular matrix. Targets are human unless otherwise specified.

**Table S 2.** List of antibodies used for immunostaining (IF), flow cytometry analysis (FCM) and immunohistochemistry (IHC-P).
| Antibody | Target | Application | Source | Identifier | Clone | Ab |
| --- | --- | --- | --- | --- | --- | --- |
| Albumin | Hepatocytes | FCM (1:100) | Dako | A0001 | Rb-Pc | 1° |
| Albumin | Hepatocytes | IF (1:100) | Bethyl | A80-229A | Gt-Pc | 1° |
| AFP | Hepatocytes | IF (1:300) | Dako | A0008 | Rb-Pc | 1° |
| KRT19 | Cholangiocytes | IF/IHC-P (1:100) | AbCam | ab7754 | Ms-Mc | 1° |
| KRT19 | Cholangiocytes | IF/IHC-P (1:100) | AbCam | ab76539 | Rb-Mc | 1° |
| KRT7 | Cholangiocytes | IF (1:100) | AbCam | ab68459 | Rb-Mc | 1° |
| KRT7 | Cholangiocytes | IF (1:100) | AbCam | ab216016 | Ms-Mc | 1° |
| SOX9 | Cholangiocytes | IF (1:100) | R&D | AF3075 | Gt-Pc | 1° |
| SOX9 | Cholangiocytes | IF (1:100) | AbCam | ab185230 | Rb-Mc | 1° |
| ITGβ1 | Integrin | IF (1:100) | AbCam | ab30394 | Ms-Mc | 1° |
| Active ITGβ1 | Integrin | IF (1:100) | AbCam | ab30394 | Ms-Mc | 1° |
| ITGα2 | Integrin | IF (1:100) | Biotechne | MAB12332 | Rt-Mc | 1° |
| ITGαV | Integrin | IHC-P (1:100) | AbCam | ab179475 | Rb-Mc | 1° |
| ITGα2β1 | Integrin | IF (1:100) | AbCam | ab230290 | Ms-Mc | 1° |
| ITGβ1 (AIIB2) | Integrin | Blocking | Thermo | MABT409 | Rt-Mc | 1° |
| Active YAP1 | YAP/TAZ | IF/IHC-P (1:100) | AbCam | ab205270 | Rb-Mc | 1° |
| VIM | Non-parenchymal | IF/IHC-P (1:100) | AbCam | ab92547 | Rb-Mc | 1° |
| COL IV | ECM | IHC-P (1:500) | AbCam | ab6586 | Rb-Pc | 1° |
| COL I | ECM | IHC-P (1:1,000) | AbCam | ab138492 | Rb-Mc | 1° |
| Fibronectin | ECM | IHC-P (1:100) | AbCam | ab2413 | Rb-Pc | 1° |
| LAMA5 | ECM | IHC-P (1:100) | AbCam | ab77175 | Ms-Mc | 1° |
| Laminin | ECM | IHC-P (1:100) | AbCam | ab11575 | Rb-Pc | 1° |
| CD31 | LSECs | IHC-P (1:100) | AbCam | ab9498 | Ms-Mc | 1° |
| αRb 488 | 1° Ab | IF/FCM/IHC-P (1:1,000) | Invitro | A21206 | Dk-Pc | 2° |
| αRb 568 | 1° Ab | IF/IHC-P (1:1,000) | Invitro | A10042 | Dk-Pc | 2° |
| αRb 647 | 1° Ab | IF/IHC-P (1:1,000) | Invitro | A31573 | Dk-Pc | 2° |
| αMs 488 | 1° Ab | IF/IHC-P (1:1,000) | Invitro | A21202 | Dk-Pc | 2° |
| αMs 568 | 1° Ab | IF/IHC-P (1:1,000) | Invitro | A10037 | Dk-Pc | 2° |
| αMs 647 | 1° Ab | IF/IHC-P (1:1,000) | Invitro | A31571 | Dk-Pc | 2° |
| αGt 488 | 1° Ab | IF/IHC-P (1:1,000) | Invitro | A11055 | Dk-Pc | 2° |
| αGt 568 | 1° Ab | IF/IHC-P (1:1,000) | Invitro | A11057 | Dk-Pc | 2° |
| αGt 647 | 1° Ab | IF/IHC-P (1:1,000) | Invitro | A21447 | Dk-Pc | 2° |
| αRt 488 | 1° Ab | IF/IHC-P (1:1,000) | Invitro | A21208 | Dk-Pc | 2° |
| αRt 568 | 1° Ab | IF/IHC-P (1:1,000) | Invitro | A78946 | Dk-Pc | 2° |
| αRt 647 | 1° Ab | IF/IHC-P (1:1,000) | Invitro | A78947 | Dk-Pc | 2° |

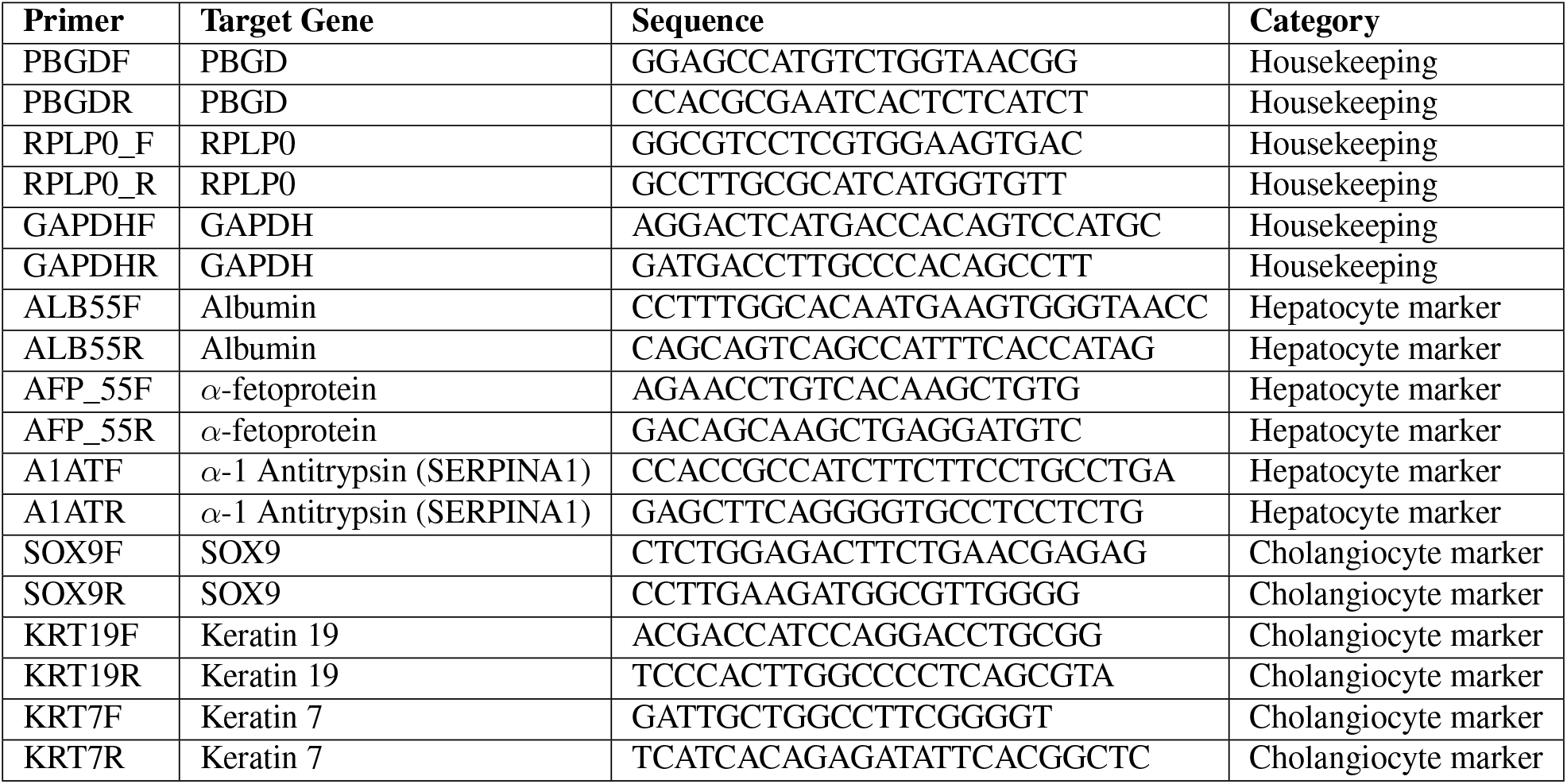
Primers used in real-time quantitative PCR analysis.

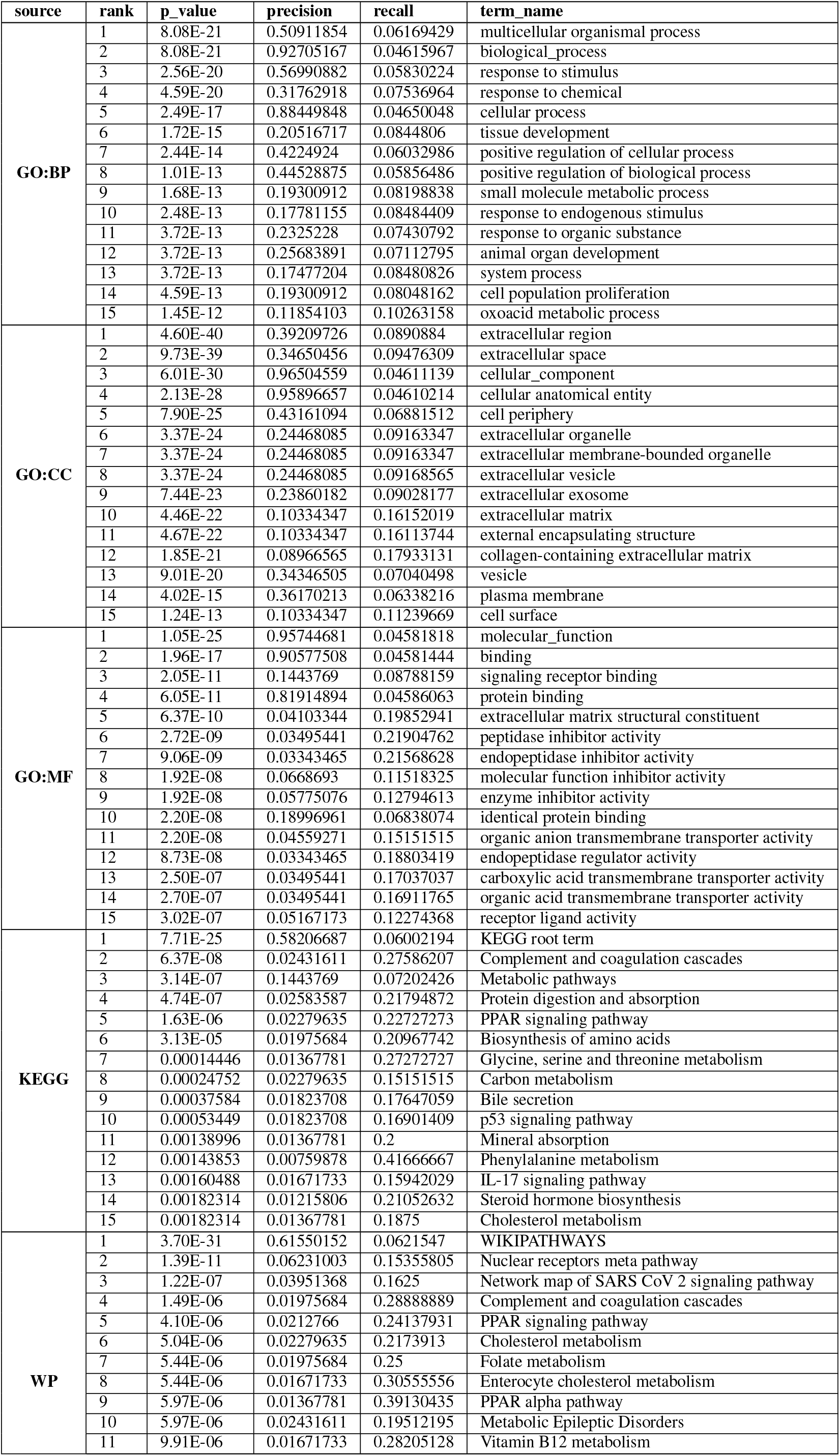
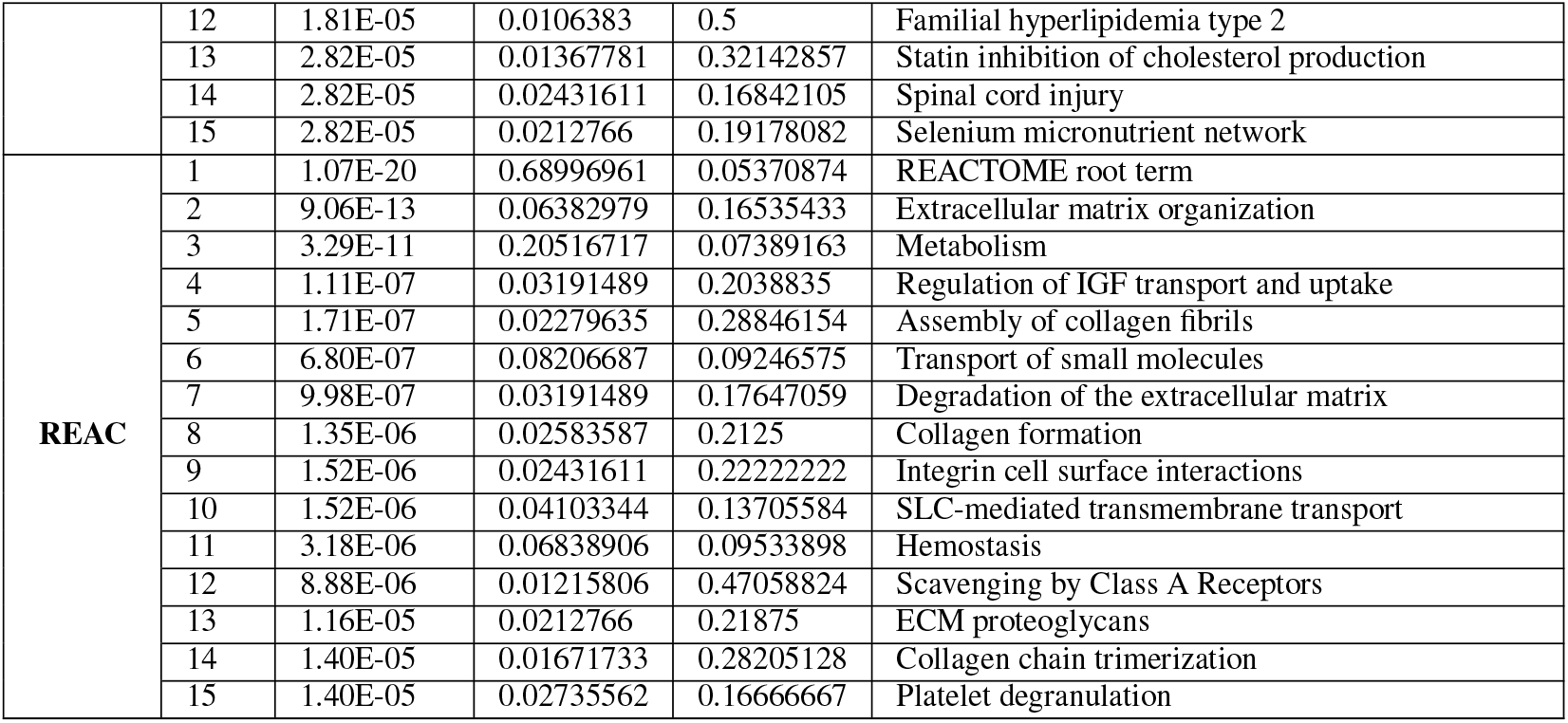
GSEA analysis of DE genes between day 16 CLCs on StemBond (0.7kPa and 3.5kPa) and ICO Controls.

GSEA analysis of DE genes between day 26 CLCs on StemBond (0.7kPa and 3.5kPa) and ICO Controls.

**Table S 5.** GSEA analysis of DE genes between day 26 CLCs on StemBond (0.7kPa and 3.5kPa) and ICO Controls.
| source | rank | p_value | precision | recall | term_name |
| --- | --- | --- | --- | --- | --- |
| GO:BP | 1 | 4.89E-23 | 0.92696629 | 0.05003032 | biological_process |
|  | 2 | 6.27E-23 | 0.57303371 | 0.06351183 | response to stimulus |
|  | 3 | 2.86E-21 | 0.50140449 | 0.06595234 | multicellular organismal process |
|  | 4 | 5.24E-21 | 0.31601124 | 0.08052971 | response to chemical |
|  | 5 | 4.71E-19 | 0.88342697 | 0.05038449 | cellular process |
|  | 6 | 8.44E-19 | 0.13764045 | 0.12189055 | oxoacid metabolic process |
|  | 7 | 9.98E-19 | 0.20926966 | 0.09418458 | small molecule metabolic process |
|  | 8 | 9.98E-19 | 0.13764045 | 0.12113721 | organic acid metabolic process |
|  | 9 | 1.78E-17 | 0.13202247 | 0.11944091 | carboxylic acid metabolic process |
|  | 10 | 2.66E-14 | 0.19522472 | 0.0870382 | tissue development |
|  | 11 | 4.41E-13 | 0.68539326 | 0.05344431 | biological regulation |
|  | 12 | 1.29E-12 | 0.25140449 | 0.07461442 | animal organ development |
|  | 13 | 2.00E-12 | 0.42696629 | 0.06162579 | developmental process |
|  | 14 | 2.72E-12 | 0.08848315 | 0.126 | small molecule biosynthetic process |
|  | 15 | 3.37E-12 | 0.4002809 | 0.0626236 | anatomical structure development |
| GO:CC | 1 | 1.53E-38 | 0.37780899 | 0.09301521 | extracellular region |
|  | 2 | 5.73E-37 | 0.33146067 | 0.09862098 | extracellular space |
|  | 3 | 1.10E-29 | 0.95926966 | 0.04965467 | cellular_component |
|  | 4 | 1.98E-28 | 0.95365169 | 0.04966718 | cellular anatomical entity |
|  | 5 | 3.28E-25 | 0.24157303 | 0.09789414 | extracellular organelle |
|  | 6 | 3.28E-25 | 0.24157303 | 0.09789414 | extracellular membrane-bounded organelle |
|  | 7 | 3.28E-25 | 0.24157303 | 0.09794989 | extracellular vesicle |
|  | 8 | 1.82E-23 | 0.23455056 | 0.0960322 | extracellular exosome |
|  | 9 | 3.73E-23 | 0.41432584 | 0.07198633 | cell periphery |
|  | 10 | 2.26E-20 | 0.33848315 | 0.07545398 | vesicle |
|  | 11 | 3.61E-19 | 0.08146067 | 0.17791411 | collagen-containing extracellular matrix |
|  | 12 | 6.35E-19 | 0.09269663 | 0.15714286 | extracellular matrix |
|  | 13 | 6.69E-19 | 0.09269663 | 0.1567696 | external encapsulating structure |
|  | 14 | 1.77E-15 | 0.35533708 | 0.06791946 | plasma membrane |
|  | 15 | 1.10E-13 | 0.06320225 | 0.16605166 | endoplasmic reticulum lumen |
| GO:MF | 1 | 2.57E-26 | 0.95365169 | 0.04942136 | molecular_function |
|  | 2 | 2.08E-18 | 0.90308989 | 0.04949962 | binding |
|  | 3 | 7.67E-13 | 0.82303371 | 0.04991057 | protein binding |
|  | 4 | 6.07E-09 | 0.13061798 | 0.08683473 | signaling receptor binding |
|  | 5 | 2.38E-08 | 0.03651685 | 0.19117647 | extracellular matrix structural constituent |
|  | 6 | 7.39E-08 | 0.18398876 | 0.07229581 | identical protein binding |
|  | 7 | 9.81E-08 | 0.03370787 | 0.19047619 | carboxylic acid transmembrane transporter activity |
|  | 8 | 1.02E-07 | 0.03370787 | 0.18897638 | organic acid transmembrane transporter activity |
|  | 9 | 2.26E-07 | 0.04073034 | 0.15425532 | organic anion transmembrane transporter activity |
|  | 10 | 3.07E-06 | 0.04775281 | 0.12318841 | receptor ligand activity |
|  | 11 | 3.13E-06 | 0.08005618 | 0.09223301 | oxidoreductase activity |
|  | 12 | 3.14E-06 | 0.10533708 | 0.08169935 | transporter activity |
|  | 13 | 3.70E-06 | 0.04775281 | 0.12099644 | signaling receptor activator activity |
|  | 14 | 6.98E-06 | 0.0491573 | 0.11551155 | signaling receptor regulator activity |
|  | 15 | 8.26E-06 | 0.02668539 | 0.17924528 | peptidase inhibitor activity |
| KEGG | 1 | 4.87E-28 | 0.56882022 | 0.06627393 | KEGG root term |
|  | 2 | 6.66E-10 | 0.15308989 | 0.08288973 | Metabolic pathways |
|  | 3 | 1.34E-07 | 0.02247191 | 0.27586207 | Complement and coagulation cascades |
|  | 4 | 2.20E-07 | 0.0252809 | 0.23076923 | Protein digestion and absorption |
|  | 5 | 2.34E-07 | 0.02247191 | 0.25806452 | Biosynthesis of amino acids |
|  | 6 | 3.83E-06 | 0.02106742 | 0.22727273 | PPAR signaling pathway |
|  | 7 | 6.49E-06 | 0.0252809 | 0.18181818 | Carbon metabolism |
|  | 8 | 0.00024098 | 0.01264045 | 0.27272727 | Glycine, serine and threonine metabolism |
|  | 9 | 0.00034311 | 0.01825843 | 0.17808219 | ECM-receptor interaction |
|  | 10 | 0.00043251 | 0.01404494 | 0.22222222 | Mineral absorption |
|  | 11 | 0.00061906 | 0.01685393 | 0.17647059 | Bile secretion |
|  | 12 | 0.00061906 | 0.01685393 | 0.17647059 | IL-17 signaling pathway |
|  | 13 | 0.0007729 | 0.01685393 | 0.17142857 | p53 signaling pathway |
|  | 14 | 0.00092433 | 0.01264045 | 0.2195122 | Bladder cancer |
|  | 15 | 0.00136112 | 0.01123596 | 0.23529412 | Fat digestion and absorption |
| WP | 1 | 2.22E-33 | 0.62640449 | 0.06633943 | WIKIPATHWAYS |
|  | 2 | 4.29E-11 | 0.05898876 | 0.15730337 | Nuclear receptors meta-pathway |
|  | 3 | 3.31E-07 | 0.0238764 | 0.24637681 | Cholesterol metabolism |
|  | 4 | 3.31E-07 | 0.01966292 | 0.31111111 | Complement and coagulation cascades |
|  | 5 | 3.31E-07 | 0.02106742 | 0.28846154 | Folate metabolism |
|  | 6 | 3.31E-07 | 0.03651685 | 0.1625 | Network map of SARS-CoV-2 signaling pathway |
|  | 7 | 3.96E-07 | 0.04634831 | 0.13360324 | IL-24 Signaling pathway |
|  | 8 | 1.11E-06 | 0.01685393 | 0.33333333 | Enterocyte cholesterol metabolism |
|  | 9 | 2.71E-06 | 0.01123596 | 0.53333333 | Blood clotting cascade |
|  | 10 | 2.71E-06 | 0.02247191 | 0.21917808 | Selenium micronutrient network |
|  | 11 | 2.71E-06 | 0.0238764 | 0.20731707 | Metabolic Epileptic Disorders |
|  | 12 | 4.68E-06 | 0.01966292 | 0.24137931 | PPAR signaling pathway |
|  | 13 | 5.30E-06 | 0.01404494 | 0.35714286 | Statin inhibition of cholesterol production |
|  | 14 | 8.61E-06 | 0.01264045 | 0.39130435 | PPAR-alpha pathway |
|  | 15 | 1.35E-05 | 0.05477528 | 0.10209424 | Pleural mesothelioma |
| REAC | 1 | 1.94E-18 | 0.67275281 | 0.05670652 | REACTOME root term |
|  | 2 | 8.94E-13 | 0.20786517 | 0.08105148 | Metabolism |
|  | 3 | 9.16E-10 | 0.05477528 | 0.15354331 | Extracellular matrix organization |
|  | 4 | 3.49E-07 | 0.08146067 | 0.09931507 | Transport of small molecules |
|  | 5 | 3.78E-07 | 0.02949438 | 0.2038835 | Regulation of IGF transport and uptake |
|  | 6 | 3.42E-06 | 0.03932584 | 0.14213198 | SLC-mediated transmembrane transport |
|  | 7 | 3.42E-06 | 0.01966292 | 0.26923077 | Assembly of collagen fibrils |
|  | 8 | 4.45E-06 | 0.0238764 | 0.2125 | Collagen formation |
|  | 9 | 5.71E-06 | 0.06601124 | 0.09957627 | Hemostasis |
|  | 10 | 1.42E-05 | 0.02808989 | 0.16806723 | Degradation of the extracellular matrix |
|  | 11 | 1.84E-05 | 0.01123596 | 0.47058824 | Scavenging by Class A Receptors |
|  | 12 | 2.35E-05 | 0.01404494 | 0.33333333 | Amino acid transport across the plasma membrane |
|  | 13 | 3.66E-05 | 0.01544944 | 0.28205128 | Collagen chain trimerization |
|  | 14 | 3.66E-05 | 0.01825843 | 0.23214286 | Non-integrin membrane-ECM interactions |
|  | 15 | 7.05E-05 | 0.01404494 | 0.29411765 | Binding and Uptake of Ligands by Scavenger Receptors |

## Notes

### Competing Interest Statement

The authors have declared no competing interest.

## References

Brevini, Teresa, Olivia C. Tysoe, and Fotios Sampaziotis (2020). Tissue engineering of the biliary tract and modelling of cholestatic disorders.

Roos, Floris J.M. et al. (2022). “Human branching cholangiocyte organoids recapitulate functional bile duct formation”. In: Cell Stem Cell 29.5.

Maroni, Luca et al. (2015). Functional and Structural Features of Cholangiocytes in Health and Disease.

Boyer j. L., and J. R. Bloomer (1974). “Canalicular bile secretion in man. Studies utilizing the biliary clearance of [14C]mannitol”. In: Journal of Clinical Investigation 54.4.

Sampaziotis, Fotios, Daniele Muraro, et al. (Feb. 2021). “Cholangiocyte organoids can repair bile ducts after transplantation in the human liver”. In: Science 371.6531, pp. 839–846.

Dhawan, Anil et al. (2020). “Alginate microencapsulated human hepatocytes for the treatment of acute liver failure in children”. In: Journal of Hepatology 72.5.

Sampaziotis, Fotios, Charis-Patricia Segeritz, and Ludovic Vallier (July 2015). “Potential of human induced pluripotent stem cells in studies of liver disease”. In: Hepatology 62.1, pp. 303–311.

Sampaziotis, Fotios, Miguel de Brito, et al. (Aug. 2015). “Cholangiocytes derived from human induced pluripotent stem cells for disease modeling and drug validation”. en. In: Nature Biotechnology 33.8, pp. 845–852.

Sampaziotis, Fotios, Miguel Cardoso De Brito, et al. (Apr. 2017). “Directed differentiation of human induced pluripotent stem cells into functional cholangiocyte-like cells”. In: Nature Protocols 12.4, pp. 814–827.

Ogawa, Mina, Shinichiro Ogawa, et al. (July 2015). “Directed differentiation of cholangiocytes from human pluripotent stem cells”. In: Nature Biotechnology 2015 33:8 33.8, pp. 853–861.

Ogawa, Mina, Jia Xin Jiang, et al. (Nov. 2021). “Generation of functional ciliated cholangiocytes from human pluripotent stem cells”. In: Nature Communications 2021 12:1 12.1, pp. 1–19.

Dianat, Noushin et al. (2014). “Generation of functional cholangiocyte-like cells from human pluripotent stem cells and HepaRG cells”. In: Hepatology (Baltimore, Md.) 60.2, p. 700.

De Assuncao, Thiago M. et al. (June 2015). “Development and characterization of human-induced pluripotent stem cell-derived cholangiocytes”. In: Laboratory Investigation 95.6, pp. 684–696.

Matsui, Satoshi et al. (Mar. 2019). “Differentiation and isolation of iPSC-derived remodeling ductal plate-like cells by use of an AQP1-GFP reporter human iPSC line”. In: Stem cell research 35.

Fiorotto, Romina et al. (Mar. 2018). “Src kinase inhibition reduces inflammatory and cytoskeletal changes in ?F508 human cholangiocytes and improves CFTR correctors efficacy”. In: Hepatology (Baltimore, Md.) 67.3, p. 972.

Hannan, Nicholas R.F., Charis Patricia Segeritz, et al. (Feb. 2013). “Production of hepatocyte-like cells from human pluripotent stem cells”. In: Nature Protocols 8.2, pp. 430–437.

Gordillo, Miriam, Todd Evans, and Valerie Gouon-Evans (June 2015). Orchestrating liver development.

Ober, Elke A and Frédéric P Lemaigre (May 2018). “Development of the liver: Insights into organ and tissue morphogenesis”. English. In: Journal of Hepatology 68.5, pp. 1049–1062.

Si-Tayeb, Karim, Frédéric P. Lemaigre, and Stephen A. Duncan (2010). Organogenesis and Development of the Liver.

Kaylan, Kerim B et al. (Dec. 2018). “Spatial patterning of liver progenitor cell differentiation mediated by cellular contractility and Notch signaling”. In: eLife 7. Ed. by Gordana Vunjak-Novakovic and Didier Y Stainier, e38536.

Li, Li et al. (Mar. 2018). “Three-dimensional hepatocyte culture system for the study of Echinococcus multilocularis larval development”. en. In: PLOS Neglected Tropical Diseases 12.3, e0006309.

Pauklin, Siim and Ludovic Vallier (Feb. 2015). Activin/nodal signalling in stem cells.

Clotman, Frédéric, Patrick Jacquemin, et al. (Aug. 2005). “Control of liver cell fate decision by a gradient of TGFβ signaling modulated by Onecut transcription factors”. In: Genes and Development 19.16, pp. 1849–1854.

Clotman, Frédéric and Frédéric P. Lemaigre (Jan. 2006). Control of hepatic differentiation by activin/TGFβ signaling. Decaens,

Thomas et al. (Jan. 2008). “Stabilization of beta-catenin affects mouse embryonic liver growth and hepatoblast fate”. eng. In: Hepatology (Baltimore, Md.) 47.1, pp. 247–258.

Tanimizu, Naoki and Atsushi Miyajima (2004). Notch signaling controls hepatoblast differentiation by altering the expression of liver-enriched transcription factors.

Kodama, Yuzo et al. (2004). “The role of notch signaling in the development of intrahepatic bile ducts”. In: Gastroenterology 127.6.

Zong, Yiwei et al. (2009). “Notch signaling controls liver development by regulating biliary differentiation”. In: Development 136.10.

Jović, Marko et al. (Aug. 2018). “Distribution of Collagen I, III, and IV and Laminin in the Human Liver during Prenatal Development”. In: Cells Tissues Organs 205.3, pp. 164–177.

Segel, Michael et al. (Sept. 2019). “Niche stiffness underlies the ageing of central nervous system progenitor cells”. en. In: Nature 573.7772, pp. 130–134.

Ge, Yejing et al. (Mar. 2020). “The aging skin microenvironment dictates stem cell behavior”. In: Proceedings of the National Academy of Sciences of the United States of America 117.10, pp. 5339–5350.

Villeneuve, Clémentine et al. (Feb. 2024). “Mechanical forces across compartments coordinate cell shape and fate transitions to generate tissue architecture”. In: Nature Cell Biology 2024 26:2 26.2, pp. 207–218.

Koester, Janis et al. (July 2021). “Niche stiffening compromises hair follicle stem cell potential during ageing by reducing bivalent promoter accessibility”. In: Nature cell biology 23.7, pp. 771–781.

Bansaccal, Nordin et al. (Nov. 2023). “The extracellular matrix dictates regional competence for tumour initiation”. In: Nature 623.7988, pp. 828–835.

Dupont, Sirio (Apr. 2016). “Role of YAP/TAZ in cell-matrix adhesion-mediated signalling and mechanotransduction”. In: Experimental Cell Research 343.1, pp. 42–53.

Kim, Minwook, Juhoon So, and Donghun Shin (Oct. 2023). “PPARα activation promotes liver progenitor cell-mediated liver regeneration by suppressing YAP signaling in zebrafish”. In: Scientific Reports 2023 13:1 13.1, pp. 1–11.

Lee, Da Hye et al. (June 2016). “LATS-YAP/TAZ controls lineage specification by regulating TGFβ signaling and Hnf4α expression during liver development”. In: Nature Communications 2016 7:1 7.1, pp. 1–14.

Yang, Li et al. (2017). “A single-cell transcriptomic analysis reveals precise pathways and regulatory mechanisms underlying hepatoblast differentiation”. eng. In: Hepatology (Baltimore, Md.) 66.5, pp. 1387–1401.

Russell, Jacquelyn O. and Fernando D. Camargo (Jan. 2022). “Hippo signalling in the liver: role in development, regeneration and disease”. In: Nature Reviews Gastroenterology & Hepatology 2022 19:5 19.5, pp. 297–312.

Blackford, Samuel J.I. et al. (Feb. 2023). “RGD density along with substrate stiffness regulate hPSC hepatocyte functionality through YAP signalling”. In: Biomaterials 293, p. 121982.

Cozzolino, Angela Maria et al. (2016). “Modulating the Substrate Stiffness to Manipulate Differentiation of Resident Liver Stem Cells and to Improve the Differentiation State of Hepatocytes”. In: Stem Cells International 2016.

Mittal, Nikhil et al. (Sept. 2016). “Substrate Stiffness Modulates the Maturation of Human Pluripotent Stem-Cell-Derived Hepatocytes”. In: ACS Biomaterials Science and Engineering 2.9, pp. 1649–1657.

Desai, Seema S. et al. (July 2016). “Physiological ranges of matrix rigidity modulate primary mouse hepatocyte function in part through hepatocyte nuclear factor 4 alpha”. In: Hepatology 64.1, pp. 261–275.

Kourouklis, Andreas P, Kerim B Kaylan, and Gregory H Underhill (2016). “Substrate stiffness and matrix composition coordinately control the differentiation of liver progenitor cells”. eng. In: Biomaterials 99, pp. 82–94.

Monckton, Chase P. et al. (Nov. 2022). “Modulation of human iPSC-derived hepatocyte phenotype via extracellular matrix microarrays”. In: Acta Biomaterialia 153, pp. 216–230.

Lee, Soah, Alice E. Stanton, et al. (May 2019a). “Hydrogels with enhanced protein conjugation efficiency reveal stiffness-induced YAP localization in stem cells depends on biochemical cues”. In: Biomaterials 202, pp. 26–34.

Cosgrove, Brian D. et al. (Aug. 2016). “N-cadherin adhesive interactions modulate matrix mechanosensing and fate commitment of mesenchymal stem cells”. In: Nature Materials 2016 15:12 15.12, pp. 1297–1306.

Stanton, Alice E., Xinming Tong, and Fan Yang (Sept. 2019). “Extracellular matrix type modulates mechanotransduction of stem cells”. In: Acta biomaterialia 96, p. 310.

Labouesse, Céline et al. (2021). “StemBond hydrogels control the mechanical microenvironment for pluripotent stem cells”. In: Nature Communications 12.1, p. 6132.

Mulas, Carla et al. (2020). “Microfluidic platform for 3D cell culture with live imaging and clone retrieval”. In: Lab on a Chip 20.14.

Shiojiri, Nobuyoshi and Yoshinori Sugiyama (2004). “Immunolocalization of extracellular matrix components and integrins during mouse liver development”. In: Hepatology 40.2.

Yin, Chunyue et al. (May 2013). “Hepatic stellate cells in liver development, regeneration, and cancer”. In: The Journal of Clinical Investigation 123.5, pp. 1902–1910.

Han, Zuoning et al. (May 2021). “Integrin αVβ1 regulates procollagen I production through a non-canonical transforming growth factor β signaling pathway in human hepatic stellate cells”. In: Biochemical Journal 478.9, pp. 1689–1703.

Wesley, Brandon T. et al. (2022). “Single-cell atlas of human liver development reveals pathways directing hepatic cell fates”. In: Nature Cell Biology 24.10.

Jokinen, Johanna et al. (July 2004). “Integrin-mediated Cell Adhesion to Type I Collagen Fibrils”. In: Journal of Biological Chemistry 279.30, pp. 31956–31963.

Borrirukwanit, Kulrut et al. (Oct. 2014). “High threshold of β1 integrin inhibition required to block collagen I-induced membrane type-1 matrix metalloproteinase (MT1-MMP) activation of matrix metalloproteinase 2 (MMP-2)”. In: Cancer Cell International 14.1.

Hutton, Melanie L. et al. (Nov. 2010). “Helicobacter pylori Exploits Cholesterol-Rich Microdomains for Induction of NF-κBDependent Responses and Peptidoglycan Delivery in Epithelial Cells”. In: Infection and Immunity 78.11, p. 4523.

Dupont, Sirio et al. (June 2011). “Role of YAP/TAZ in mechanotransduction”. In: Nature 474.7350, pp. 179–184.

Molina, Laura M. et al. (July 2021). “Compensatory hepatic adaptation accompanies permanent absence of intrahepatic biliary network due to YAP1 loss in liver progenitors”. In: Cell reports 36.1, p. 109310.

Lee, Soah, Alice E Stanton, et al. (2019b). “Hydrogels with enhanced protein conjugation efficiency reveal stiffness-induced YAP localization in stem cells depends on biochemical cues”. In: Biomaterials 202, pp. 26–34.

Noce, Valeria et al. (Oct. 2019). “YAP integrates the regulatory Snail/HNF4α circuitry controlling epithelial/hepatocyte differentiation”. In: Cell Death & Disease 2019 10:10 10.10, pp. 1–13.

Szeto, Stephen G. et al. (Oct. 2016). “YAP/TAZ are mechanoregulators of TGF-b-smad signaling and renal fibrogenesis”. In: Journal of the American Society of Nephrology 27.10, pp. 3117–3128.

Koo, Ja Hyun et al. (Jan. 2020). “Induction of AP-1 by YAP/TAZ contributes to cell proliferation and organ growth”. In: Genes & development 34.1-2, pp. 72–86.

Lang, Christine, Lisa Conrad, and Dagmar Iber (June 2021). “Organ-Specific Branching Morphogenesis”. In: Frontiers in Cell and Developmental Biology 9, p. 671402.

Rizwan, Muhammad, Ana Fokina, et al. (Jan. 2021). “Photochemically Activated Notch Signaling Hydrogel Preferentially Differentiates Human Derived Hepatoblasts to Cholangiocytes”. In: Advanced Functional Materials 31.5, p. 2006116.

Sapudom, Jiranuwat et al. (Aug. 2023). “Collagen Fibril Orientation Instructs Fibroblast Differentiation Via Cell Contractility”. In: Advanced Science 10.22, p. 2301353.

Rizwan, Muhammad, Christopher Ling, et al. (Dec. 2022). “Viscoelastic Notch Signaling Hydrogel Induces Liver Bile Duct Organoid Growth and Morphogenesis”. In: Advanced Healthcare Materials 11.23, p. 2200880.

Smith, Quinton et al. (Jan. 2022). “Directing Cholangiocyte Morphogenesis in Natural Biomaterial Scaffolds”. In: Advanced Science 9.3.

Elci, Bilge Sen et al. (Mar. 2024). “Bioengineered Tubular Biliary Organoids”. In: Advanced Healthcare Materials 13.8, p. 2302912.

Tanimizu, Naoki, Atsushi Miyajima, and Keith E. Mostov (Apr. 2007). “Liver Progenitor Cells Develop Cholangiocyte-Type Epithelial Polarity in Three-dimensional Culture”. In: Molecular Biology of the Cell 18.4, p. 1472.

Chen, Chen et al. (June 2018). “Bioengineered bile ducts recapitulate key cholangiocyte functions”. In: Biofabrication 10.3, p. 034103.

Wu, Fenfang et al. (June 2019). “Generation of hepatobiliary organoids from human induced pluripotent stem cells”. In: Journal of hepatology 70.6, pp. 1145–1158.

Buxboim, Amnon et al. (2010). “How deeply cells feel: Methods for thin gels”. In: Journal of Physics Condensed Matter 22.19.

Tysoe, Olivia C et al. (June 2019). “Isolation and propagation of primary human cholangiocyte organoids for the generation of bioengineered biliary tissue”. en. In: Nature Protocols 14.6, pp. 1884–1925.

Hannan, Nicholas R.F., Robert P. Fordham, et al. (Oct. 2013). “Generation of multipotent foregut stem cells from human pluripotent stem cells”. In: Stem Cell Reports 1.4, pp. 293–306.

Hunter, Emma J. et al. (Oct. 2021). “Selectivity of the collagen-binding integrin inhibitors, TC-I-15 and obtustatin”. In: Toxicology and Applied Pharmacology 428, p. 115669.

Schindelin, Johannes et al. (July 2012). Fiji: An open-source platform for biological-image analysis.

Carpenter, Anne E. et al. (2006). “CellProfiler: Image analysis software for identifying and quantifying cell phenotypes”. In: Genome Biology 7.10.

Wolf, F. Alexander, Philipp Angerer, and Fabian J. Theis (2018). “SCANPY: Large-scale single-cell gene expression data analysis”. In: Genome Biology 19.1.

